# Exposure to herbivore-induced plant volatiles directly induces jasmonic acid and primes chemical defences in cotton plants

**DOI:** 10.64898/2026.04.09.717419

**Authors:** Kathrin Altermatt, Wenfeng Ye, Armelle Vallat, Luis Abdala-Roberts, Ted C.J. Turlings, Carlos Bustos-Segura

## Abstract

Plants can deploy alternative defensive strategies in response to airborne signals from damaged neighbours to prepare for incoming attack: a straightaway response up-regulating their defences (induction), or a primed state, leading to a faster/stronger defence response after herbivory. However, it is unclear which mechanisms are involved in each response. We used a monophagous and a polyphagous leafworm species to specifically dissect induction and priming effects of exposure to herbivore-induced plant volatiles (HIPVs). Exposure to HIPVs directly elevated jasmonate levels in undamaged receivers but did not induce defensive terpenoids or volatiles. However, HIPV-primed plants accumulated high levels of toxic terpenoids (e.g. gossypol) and emitted high quantities of volatile sesquiterpenes, when damaged by either species of caterpillars. This comprehensive study demonstrates that both defence induction and priming can be detected in cotton but occur as different, linked responses which are robust to herbivore identity, providing insights into a generalised plant communication strategy.

## Introduction

Plants have a wide range of defensive mechanisms that help them face herbivory, including the release of herbivore-induced plant volatiles (HIPVs). These HIPVs can repel herbivores and attract their natural enemies, but they may also serve as signals to prepare for a herbivore attack when perceived by other parts of the same plant (within-plant signalling) (Heil & Silva Bueno, 2007) or neighbouring plants (plant-plant signalling) (Morrell & Kessler, 2017). In such cases, exposure to HIPVs can directly induce plant defences (Baldwin & Schultz, 1983; Kost & Heil, 2006) but also lead to priming (Engelberth *et al*., 2004; Ton *et al*., 2007). In contrast to induction, priming requires two stimuli, a first priming stimulus (e.g., HIPVs) that prepares a defensive response that will be induced upon perception of a second stimulus (e.g., elicitors in insect oral secretions), to which a primed plant will respond faster and/or stronger compared to a plant in a non-primed state (Hilker *et al*., 2016; Martinez-Medina *et al*., 2016). However, it can be challenging to separate induction from priming in volatile-mediated plant-plant signalling, as exposure to HIPVs can elicit a (minor) induction response as well as provoking a primed state (Martinez-Medina *et al*., 2016; Waterman *et al*., 2024).

Domesticated cotton (*Gossypium hirsutum* L., Malvaceae) is an economically important crop cultivated in tropical and subtropical regions and used worldwide for fibre production. Cotton leaves contain glands with secondary metabolites, such as gossypol, that have antimicrobial, antiviral, and insecticidal effects (Stipanovic *et al*., 1988; Tian *et al*., 2016) and can also act as feeding deterrents (Meisner *et al*., 1977). Upon being damaged, cotton plants also release HIPVs (McCall *et al*., 1994; Röse & Tumlinson, 2005; Grandi *et al*., 2024), and multiple studies have shown undamaged cotton plants exposed to these HIPVs exhibit higher defences and resistance against herbivory. For example, cotton leaves exposed to volatiles of leaves infested by a pathogenic fungus increased their production of terpenoid aldehydes such as heliocides (Zeringue, 1987). Similarly, cotton plants exposed to volatiles of neighbouring plants attacked by herbivorous mites also had reduced oviposition by the same herbivores and were more attractive to predatory mites (Bruin *et al*., 1992). In the case of chewing insects, a reduction in oviposition by *Spodoptera littoralis* was shown on undamaged cotton plants exposed to HIPVs of damaged cotton plants (Zakir *et al*., 2013), and recent work with wild cotton showed that plants produced more extrafloral nectar and heliocides upon damage when they were previously exposed to HIPVs from neighbouring plants (Briones-May *et al*., 2023; Quijano-Medina *et al*., 2024). Finally, (Grandi *et al*., 2024) found that cotton plants exposed to HIPVs from plants damaged by *Spodoptera* spp. caterpillars for longer than 24 h showed higher volatile emissions, secondary metabolites, phytohormones and defence gene expression than those exposed to volatiles from control plants. The HIPV-exposed plants were also avoided in choice tests with caterpillars, in particular it was shown that only *de novo* synthetised volatiles after attack induce direct defences in neighbouring cotton plants.

While some studies have suggested that exposure to HIPVs enhance as well the induction of extra-floral nectar (Briones-May *et al*., 2023) and heliocides (Quijano-Medina *et al*., 2024) in wild cotton, priming has not yet been fully demonstrated in either wild or domesticated cotton, since previous studies could not distinguish priming from direct induction. A second crucial aspect that can complicate the detection of defensive responses in cotton is the timing of exposure and response. The induction of volatile emissions or phytohormone levels can be detected after a few hours or days, whereas the enhanced accumulation of terpene aldehydes such as gossypol takes considerably longer and mainly happens during the development of new tissues, where more and larger gossypol glands are formed (Bezemer *et al*., 2004; Eisenring *et al*., 2017; Mamin *et al*., 2023; Quijano-Medina *et al*., 2024). Thus, the timing of terpene aldehydes accumulation should be considered in experimental designs.

Another factor to consider in plant-plant signalling is that HIPV emissions can vary qualitatively and quantitatively depending on the pest species. It has been proposed that herbivores adapted to one plant species can better tolerate plant defences or even manipulate plant responses to their benefit (Whittaker & Feeny, 1971; Zarate *et al*., 2007; Xu *et al*., 2019). In this context, monophagous (specialist) and highly polyphagous (generalist) herbivore species may trigger different induced defence responses in plants. However, the effect of diet breadth on HIPV induction remains unclear (Rowen & Kaplan, 2016), although a few studies indicate that monophagous insects trigger lower HIPV release than polyphagous species. This is the case of rice and maize plants attacked by different caterpillar species (Sobhy *et al*., 2017; De Lange *et al*., 2020). Hence, testing for priming effects in response to attack by insects with contrasting characteristics (including diet breadth), allows testing variability (or consistency) in plant volatile-mediated defensive responses and provides insight into the underlying mechanisms.

The goal of this study was to investigate whether chemical defences are primed by HIPVs and if this priming effect differs between herbivore species with contrasting diet breadth. To do so, we first investigated whether caterpillar-induced HIPVs differ when cotton plants (emitters) are infested by the monophagous cotton leafworm (*Alabama argillacea,* Hübner 1823, Lepidoptera, Noctuidae), or the highly polyphagous Egyptian cotton leafworm (*Spodoptera littoralis,* Boisduval 1833, Lepidoptera, Noctuidae). We then exposed undamaged cotton plants (receivers) to volatiles from plants infested by *A. argillacea* or *S. littoralis* or to volatiles from undamaged plants to initially determine direct defence induction due to HIPVs exposure, and subsequently infested half of receiver plants with the same caterpillar species used to induce emitters to test for priming effects. We then measured the volatile emissions, phytohormone levels, terpene aldehyde levels and associated gene expression in receiver plants which allowed us to clearly differentiate between induced responses and priming upon HIPV exposure.

## Materials and Methods

### Plants

Cotton seeds (*Gossypium hirsutum* L., var STAM 59) were individually planted in plastic pots (diameter 6 cm, height 10 cm, 250 ml). Seeds were planted in RICOTER Schweizer Erde (RICOTER Erdaufbereitung AG, Aarberg, Switzerland) and supplemented with 10 ml of fertilizer solution (0.34%) using CAPITO Flüssigdünger Universal (Landi Schweiz AG, Dotzingen, Switzerland) once per week starting two weeks after sowing to account for nutrient deficiency in this soil. All plants were kept in phytotrons (GroBanks, CLF Plant Climatics, Wertingen, Germany) under controlled conditions (Light 14 h and 28°C, dark 10 h and 25°C, Luminosity 65 ± 10 μmol, R.H. 50 ± 10%) and were watered 3 times per week. Plants were used for experiments once they had three fully developed true leaves.

### Insects

The cotton leafworm *Alabama argillacea* (Hübner 1823, Lepidoptera, Noctuidae) was used as the monophagous herbivore*. Alabama argillacea* is present in Central and South America and is one of the main insect pests of wild and domesticated cotton in the region, feeding on plants from the tribe Gossypieae (Montandon *et al*., 1986; Silva *et al*., 2011). Larvae of *A. argillacea* from northern Yucatán, México were collected and reared to establish a colony at the University of Neuchâtel (FOEN permit: A192632-07). Adults were kept in a cage supplemented with water, apple slices and honey to feed on, and with cotton plants to lay eggs on. Leaves with eggs were removed from the cage and placed in plastic containers. Caterpillars were kept in plastic containers and fresh cotton leaves were added every other day until pupation.

The Egyptian cotton leafworm *Spodoptera littoralis* (Boisduval 1833, Lepidoptera, Noctuidae) is widely distributed in Africa, Mediterranean Europe and the Middle East (Wu *et al*., 2022). It is a highly polyphagous pest feeding on a wide variety of plants, including many crops such as tomatoes, peppers, maize and cotton, and is one of the main pest of cotton in Egypt (Salama *et al*., 1971; Wu *et al*., 2022). *Spodoptera littoralis* eggs were sent once per week from Syngenta (Syngenta AG, Basel, Switzerland). The caterpillars were fed on artificial diet based on wheat-germ (Frontier Scientific Services, Newark, United States). Before being used in experiments, *S. littoralis* caterpillars were fed on cotton leaves for approximately 24 h and starved overnight. Both insects were kept under similar conditions (L16:D8, 26 ± 2°C, 50 ± 10% R.H.).

### Plant-plant signalling experimental setup

Emitter cotton plants at 3^rd^ leaf stage (3^rd^ leaf almost fully developed) were left undamaged as control plants (emitting constitutive plant volatiles, CPVs) or infested with either two 2^nd^ instar caterpillars of *A. argillacea* or three to five 2^nd^ instar caterpillars of *S. littoralis* (emitting herbivore-induced plant volatiles, HIPVs). The number of caterpillars was chosen to ensure a similar quantity of damage on the plants. The plant pots were covered with aluminium foil and carefully placed in custom-made glass bottles (VTQ SA, Switzerland). Each bottle with these emitter plants was then connected to another bottle containing an undamaged receiver cotton plant. Filtered air was pushed in with 0.9 l/min (± 0.01 l/min) at the bottom of the bottle containing the emitter plant, passing by the top of both bottles through the connection tube and leaving the system at the bottom of the bottle containing the receiver plant (Figure S1). The receiver plants were exposed to the volatiles of the emitter plants for 72 h.

After 72 h, the bottles with the emitter and receiver plants were separated, the volatiles of the emitter plants were collected and then their leaves were cut off at the base, weighed and photographed. Receiver plants were either left undamaged (controls) or infested with one 2^nd^ instar caterpillar of either *A. argillacea* or *S. littoralis* using a clip cage on the 1^st^ leaf. Damaged receiver plants were damaged by the same species that infested their emitter couple. This resulted in a total of seven treatment groups: one group of CPV-exposed undamaged receivers, two groups of HIPV-exposed undamaged receivers, two groups of CPV-exposed damaged receivers, and two groups of HIPV-exposed damaged receivers (n = 6 per treatment).

To measure receiver gene expression and phytohormones (n = 6 per treatment) after herbivory, caterpillars were removed after 24h of damage, the 3^rd^ leaf was cut off at the base, weighed and directly frozen using liquid nitrogen. Samples were stored at −80 until further analyses. The 1^st^ leaf was cut off and photographed to assess damage.

To measure receiver volatile emissions, terpene aldehyde content and herbivore performance (n = 6 per treatment), caterpillars were left to feed for 48 h on the plant. Volatiles were collected after 24 and 48 h of infestation. During the volatile collection, the caterpillar and clip cage were removed and then placed back in a different position. After the second volatile collection, caterpillars and clip cages were removed and plants were put back in phytotrons for two weeks until they had developed the 4^th^ leaf. The 4^th^ leaf was cut off at the base and frozen in liquid nitrogen to measure terpenoid aldehydes.

### Volatile collection and analyses

To collect volatiles, plant pots were covered in aluminium foil and placed in the glass bottles. Filtered air was pushed into each bottle at the bottom (0.9 ± 0.01 l/min) and sucked out at the top (0.6 ± 0.01 l/min). Volatiles were collected by placing trapping filters containing 25 mg of 80/100 mesh Hayesep-Q adsorbent (Ohio Valley Specialist Company, Marietta, United States) for 2 h. After each collection, the filters were directly eluted with 100 μl of dichloromethane and 10 μl of internal standard were added (containing 20 ng/μl of n-octane and n-nonyl acetate). Samples were kept at −80°C until further analyses.

Volatile extracts were analysed on a gas chromatograph (Agilent 7890B) coupled to a mass spectrometer (Agilent 5977B GC/MSD) in TIC mode. 1.5 μl of each sample was injected in pulsed splitless mode into an Agilent HP-5MS column (length 30 m, diameter 0.25 mm, film thickness 0.25 μm). After injection, temperature was kept at 40°C for 3 min, then increased to 100°C at a rate of 8°C/min and then to 200°C at 5°C/min, followed by a post run period at 250°C for 3 min. Helium was used as a carrier gas and kept at constant flow of 1.1 ml/min. Compounds were identified using the NIST 17 mass spectra library (U.S. Department of Commerce, Gaithersburg, United States) and an in-house library based on authentic standards.

### Gene expression

To measure gene expression, 75 ± 5 mg of frozen powdered leaf material was used per sample. RNA was isolated using the GeneJET Plant Purification Mini Kit (Thermo Fisher Scientific Baltics UAB, Vilnius, Lithuania) according to the manufacturer’s instructions and complete DNA removal was performed using the RNase-Free DNase Set (QIAGEN, Hilden, Germany). Each total RNA sample (300 ng) was reverse transcribed using the GoScript™ Reverse Transcription System (Promega). Real-time qPCR was performed on the Rotor-Gene™ 6000 (Corbett Research) platform. The qPCR mix consisted of 10 μl GoTaq® qPCR Master Mix (Promega), 8.2 μl H_2_O, 0.4 μl each primer (10 μM) and 1 μl of cDNA sample. The qPCR was performed using 50 cycles with the following temperature curve: 15 s at 95°C and 60 s at 60°C. The melt curve was obtained by ramping from 50°C to 99°C, rising by 1°C each step and wait for 5 s for each step afterwards. The *Histone* gene (GenBank accession number: AF024716) was used as the reference gene for the transcript profiling. Primers used for real-time qPCR are listed in Supplementary Table S9. The threshold cycle (Ct) per sample was calculated for each treatment. Undamaged plants were designated as calibrator, and the relative expression levels were calculated using the 2^-△△Ct^ method (Livak & Schmittgen, 2001).

### Phytohormones

Levels of abscisic acid (ABA), indole-3-acetic acid (IAA), jasmonic acid (JA), jasmonic acid-isoleucine (JA-Ile), 12-oxophytodienoic acid (OPDA), and salicylic acid (SA) were determined with liquid chromatography (LC) and mass spectrometry (MS). To do so, 80 ± 5 mg of frozen leaf powder was transferred to a 1.5 ml Eppendorf tube, 990 μl of ethylacetate : formic acid (99.5:0.5, v/v) and 10 μl of internal standard (containing isotopically labelled hormones at 100 ng/ml in water) were added, and each sample was vortexed immediately for 10 s. Five to ten glass beads (1.25 – 1.65 mm diameter; Sigmund Lindner GmbH, Warmensteinach, Germany) were added, the tubes were placed in a mixer mill (Tissue Lyser II, Qiagen) at 30 Hz for 3 min and then centrifuged (PrismTM Microcentrifuge, Labnet) at 14’000 g for 3 min. The supernatant was then transferred to a 2 ml Eppendorf tube. The remaining pellet was re-extracted with 500 μl of ethylacetate : formic acid (99.5:0.5, v/v), centrifuged for 3 min at 14’000 g and the two supernatants were combined. Using a centrifugal evaporator (CentriVap®, Labconco) at 35°C, the samples were evaporated for 60 min. 200 μl of MeOH 50% was added to the residue and the tube was placed for 2 min in an ultrasonic bath and vortexed for 10 s to resuspend the residue. The sample was centrifuged for 3 min at 14’000 g and 120 μl of the supernatant was transferred in a 2 ml glass vial (Interchim, Montluçon, France) with a 250 μl insert (Interchim, Montluçon, France). Samples were analysed using Ultra High Performance Liquid Chromatography (UPLC Acquity Waters) coupled with an MS-MS (ABSciex QTrap 6500+), using the settings described in (Glauser *et al*., 2014).

### Terpene aldehydes

To extract terpene aldehydes, 50 ± 5 mg of powdered leaf were added to a 1.5 ml Eppendorf tube and 200 μl of acetonitrile : milliQwater : formic acid (80 : 18.5 : 1.5, v/v) was added. Each sample was placed in an ultrasonic bath for 5 min and centrifuged for 3 min at 8’000 g. 100 μl of the supernatant was transferred in a 2 ml glass vial with a 250 μl insert. Samples were analysed using Ultra High Performance Liquid Chromatography (Ultimate 3000 Dionex, Thermo Fisher Scientific, MA, USA) coupled to a DAD detector set at 288 ± 2 nm (Ultimate 3000 Dionex, Thermo Fisher Scientific, MA, USA). A 10 μl aliquot of each sample was injected onto an ACQUITY UPLC® BEH C18 column (2.1 x 100mm, 1.7μm) (Waters, MA, USA). The flow rate was held constant at 0.45 ml/min and the temperature was kept at 40°C. The mobile phase A consisted of 0.05% formic acid in MilliQ water (18 Ω) and the mobile phase B of 0.05% formic acid in acetonitrile (HiPerSolv, VWR Chemicals®, France). The following gradient was used: 45-90% B in 8 min, 90-100% B in 0.5 min, holding at 100% for 2.5 min followed by re-equilibration at 45% B for 3.5 min. Gossypol, hemigossypolone and heliocides (grouped H1 to H3) were identified by their retention time. Quantification was done by calculating a calibration equation obtained by linear regression from six calibration points (5 to 250 μg/ml) in gossypol equivalents.

### Statistical analyses

Statistical tests were carried out in R (version 4.4.2) (R Core Team, 2024).

For volatile emissions from emitter plants, generalised linear models with a gamma error distribution (logarithmic link function) were used. Differences among treatments (Undamaged, *A. argillacea*, *S. littoralis*) were tested with Wald chi-square tests (*car* package) (Fox & Weisberg, 2019), and the p-values for all individual compounds were corrected (false discovery rate correction). Post-hoc tests were performed as tukey-adjusted pairwise comparisons on estimated marginal means from the *emmeans* package (Lenth *et al*., 2025). Leaf damage in emitter plants was compared using a linear model from the package *lme4* (Bates *et al*., 2015).

For receiver plants, a two-model approach was used to test for differences among treatments in volatile emissions, leaf damage, phytohormone levels, gene expression and terpene aldehyde levels. To test the direct effect volatile exposure on defence levels (induction), only the undamaged receiver plants were included, and just three factors (CPVs-undamaged, HIPVs-undamaged from *A. argillacea*, HIPVs-undamaged from *S. littoralis*) were included in one model. To test the priming effect, the undamaged receiver plants exposed to CPVs (CPVs-undamaged) were removed from the main dataset, and the model tested the effects of treatment (three levels: HIPVs-undamaged, CPVs-damaged, HIPVs-damaged) and insect species (two levels: *A. argillacea* or *S. littoralis*). Depending on the distribution, generalised linear models with a gamma error distribution (logarithmic link function) or linear models from the package *lme4* were used (Bates *et al*., 2015). GLMs were followed by Wald chi-square tests, and LMs were followed by F-tests (*car* package) to test for the significance of the effect of each factor (treatment, insect and their interaction) (Fox & Weisberg, 2019). Post-hoc tests were performed as pairwise comparisons on estimated marginal means from the *emmeans* package (Lenth *et al*., 2025).

For both emitter and receiver plants, volatile blend composition was compared using redundancy analysis (RDA) from the *vegan* package on centre log-ratio transformed data, and the model was tested with a test with 999 permutations (Oksanen *et al*., 2025). Pairwise comparisons among the treatments were tested with 999 permutations and the p values adjusted (Holm method). To link volatile emissions in emitters with phytohormone and terpene aldehyde levels in receivers, we used sparse partial least squares (sPLS) models and the network function from the *mixOmics* package (Rohart *et al*., 2017). All other figures were created using the packages *ggpubr* and *ggplot2* (Wickham, 2016; Kassambara, 2025).

## Results

### Emitter Plants

Volatile emission of cotton plants differed among damage treatments (χ^2^_(2)_ = 134.44, p < 0.0001, Figure 1A). Cotton plants attacked by *A. argillacea* or *S. littoralis* caterpillars emitted significantly more volatiles than undamaged plants after 72 h of infestation, with no difference between the herbivore species in overall emissions (Figure 1A). According to a multivariate RDA model, the volatile blend composition was also different among damage treatments (F_(2)_ = 11.789, p = 0.001), and pairwise comparisons showed significant differences between the volatile blend composition of undamaged plants and the plants attacked by *A. argillacea* or *S. littoralis*, with again no significant difference between the herbivore species (Undamaged vs. *A. argillacea*: F_(1)_ = 23.166, p = 0.003; Undamaged vs. *S. littoralis*: F_(1)_ = 14.868, p = 0.003; *A. argillacea* vs. *S. littoralis*: F_(1)_ = 1.8746, p = 0.071, Figure 1B). However, when looking at individual compounds, plants attacked by *A. argillacea* emitted significantly more benzoic acid and methyl salicylate than plant attacked by *S. littoralis*, whereas plants attacked by *S. littoralis* released significantly more decyne, pentadecane, phenyl methanol, γ-bisabolene, and caryophyllene oxide (Figure 1C, Supplementary Table S1). There was no difference in the average leaf area consumed by *A. argillacea* and *S. littoralis* (F_(1)_ = 2.2751, p = 0.14, Supplementary Figure S2A).

**Figure 1.**
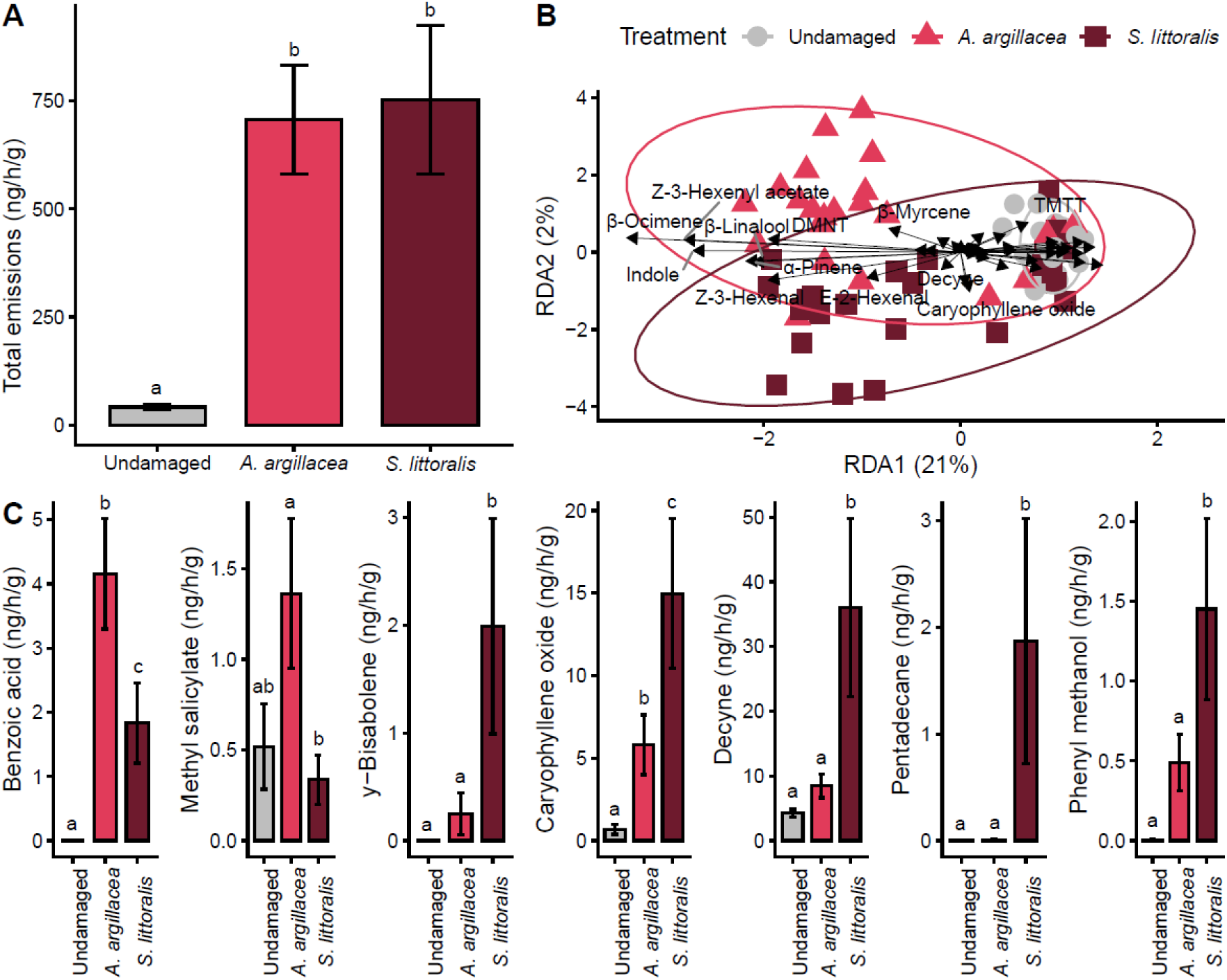
Volatile emissions of differently treated emitter plants (n = 24). Plants were either left undamaged or infested for 72 h by *A. argillacea* or *S. littoralis*, after which volatiles were collected for 2 h. **(A)** Mean overall emission rate (± SE) in ng per hour per g of fresh aboveground mass. **(B)** Redundancy analysis (RDA) with ellipses indicating the 95% confidence interval. Arrows are scaled by the factor 2.5 for clarity. **(C)** Mean emission rate (± SE) in ng per hour per g of fresh aboveground mass of all compounds that were significantly different between plants damaged by *A. argillacea* and *S. littoralis* according to univariate models (full results in Table S1). Different letters indicate significant differences (Tukey-adjusted pairwise comparisons).

### Receiver plants

To test for direct induction of volatiles in receiver plants, we first compared only undamaged receiver plants that had been exposed for 72 h. We found no significant differences in emissions between plants exposed to herbivore-induced plant volatiles (HIPVs, from damaged emitter plants) from either insect species or constitutive plant volatiles (CPVs, from undamaged emitter plants) collected either 24 h or 48 h after exposure (24h: χ^2^_(1)_ = 0.0109, p = 0.91; 48 h: χ^2^_(1)_ = 0.2596, p = 0.61, Figure 2A, Supplementary Table S2).

**Figure 2.**
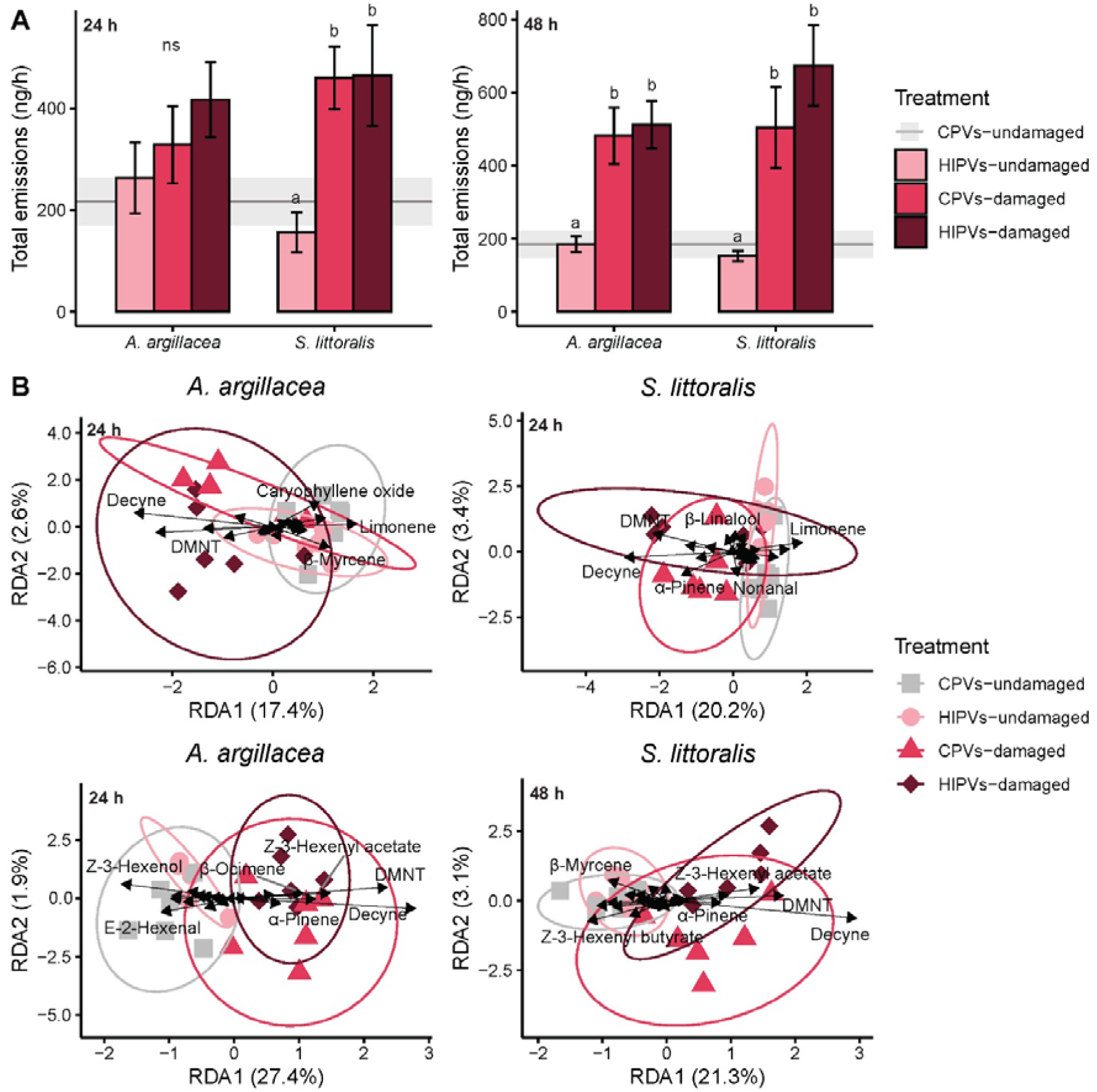
Total volatile emissions of differently treated receiver plants (n = 12). Receiver plants in 3^rd^ leaf stage were first exposed for 72 h to herbivore-induced plant volatiles from damaged emitter plants *(HIPVs*) or constitutive plant volatiles from undamaged emitter plants (*CPVs*), and were subsequently subjected to herbivory themselves (*damaged*) on the 1^st^ leaf for 48 h or left undamaged (*undamaged*). **(A)** Mean emission rate (± SE) in ng per hour. Different letters indicate significant differences (Tukey-adjusted pairwise comparisons). No differences were found between *HIPVs-undamaged* and *CPVs-undamaged* plants (baseline). **(B)** Redundancy analysis (RDA) of total emissions with ellipses indicating the 95% confidence interval. Arrows are scaled by a factor 2.5 for clarity.

To test for priming effects on volatiles emissions we next tested the effect of treatment (exposure to HIPVs and/or insect damage) in receiver plants, herbivore species and their interaction. After 24 h of volatile exposure, we found that plant treatment had an overall effect on volatile emissions in receiver plants but we did not detect a significant effect of insect species or interaction (Treatment χ^2^_(2)_ = 12.435, p = 0.002, Insect χ^2^_(1)_ = 0.0172, p = 0.9, Treatment:Insect χ^2^_(2)_ = 4.1761, p = 0.1, Figure 2A). For *S. littoralis*, damage significantly increased volatile emissions compared to undamaged plants exposed to HIPVs, but damaged receivers exposed to HIPVs did not release more total volatiles than damaged receivers exposed to CPVs. For *A. argillacea* the patterns were similar but non-significant (Figure 2A, Supplementary Table S2). After 48 h we found similar results, except that there were no differences between herbivore treatments (Treatment χ^2^_(2)_ = 67.199, p < 0.0001, Insect χ^2^_(1)_ = 0.121, p = 0.73, Treatment:Insect χ^2^_(2)_ = 2.356, p = 0.31, Figure 2A, Supplementary Table S2). The composition of the volatile blends also showed differences among treatments, for both time points and insect species, mainly influenced by 4,8-dimethyl-1,3,7-nonatriene (DMNT) and decyne, both of which were more dominant in damaged plants according to the RDA (24 h *A. argillacea* model: F_(3)_ = 1.8329, p = 0.026; 24 h *S. littoralis* model: F_(3)_ = 2.2539, p = 0.008; 48 h *A. argillacea* model: F_(3)_ = 2.9429, p = 0.002; 48 h *S. littoralis* model: F_(3)_ = 2.2902, p = 0.002, Figure 2B, Supplementary Table S3).

An inspection of patterns of individual compounds showed a significant effect of treatment for some compounds after 24 h (β-linalool: Treatment χ^2^_(2)_ = 27.0301, p < 0.0001, Insect χ^2^_(1)_ = 0.7269, p = 0.39, Treatment:Insect χ^2^_(2)_ = 1.4831, p = 0.48; β-pinene: Treatment χ^2^_(2)_ = 13.787, p = 0.001, Insect χ^2^_(1)_ = 1.0559, p = 0.30, Treatment:Insect χ^2^_(2)_ = 2.6456, p = 0.27; DMNT: Treatment χ^2^_(2)_ = 86.469, p < 0.0001, Insect χ^2^_(1)_ = 0.0004, p = 0.95, Treatment:Insect χ^2^_(2)_ = 7.619, p = 0.022). And while there seemed to be a trend for higher emissions of these compounds by damaged receiver plants that had been exposed to HIPVs compared to the damaged receivers exposed to CPVs after 24 h, the pairwise comparisons were not statistically significant (Figure 3A, Supplementary Table S4). Yet, after 48 h, treatment had a significant effect on several HIPVs (β-linalool: Treatment χ^2^_(2)_ = 50.6481, p < 0.0001, Insect χ^2^_(1)_ = 1.623, p = 0.20, Treatment:Insect χ^2^_(2)_ = 0.154, p = 0.93; DMNT: Treatment χ^2^_(2)_ = 168.453, p < 0.0001, Insect χ^2^_(1)_ = 0.179, p = 0.67, Treatment:Insect χ^2^_(2)_ = 5.091, p = 0.78; α-farnesene: Treatment χ^2^_(2)_ = 17.365, p = 0.00017, Insect χ^2^_(1)_ = 2.6172, p = 0.11, Treatment:Insect χ^2^_(2)_ = 2.7994, p = 0.25; β-farnesene: Treatment χ^2^_(2)_ = 31.322, p < 0.0001, Insect χ^2^_(1)_ = 0.0071, p = 0.93, Treatment:Insect χ^2^_(2)_ = 0.2848, p = 0.87; γ-bisabolene: Treatment χ^2^_(2)_ = 18.0668, p = 0.00021, Insect χ^2^_(1)_ = 6.5872, p = 0.010, Treatment:Insect χ^2^_(2)_ = 14.9285, p = 0.00057). More specifically, damaged receivers exposed to HIPVs emitted higher quantities of α-farnesene and β-farnesene for *A. argillacea*, and higher quantities of β-farnesene and γ-bisabolene for S*. littoralis* (Figure 3B, Supplementary Table S4). Consumed leaf area of receiver plants was not significantly different between treatments (Treatment χ^2^_(1)_ = 1.0574, p = 0.32, Insect χ^2^_(1)_ = 1.1993, p = 0.29, Treatment:Insect χ^2^_(1)_ = 0.1695, p = 0.68, Supplementary Figure S2B).

**Figure 3.**
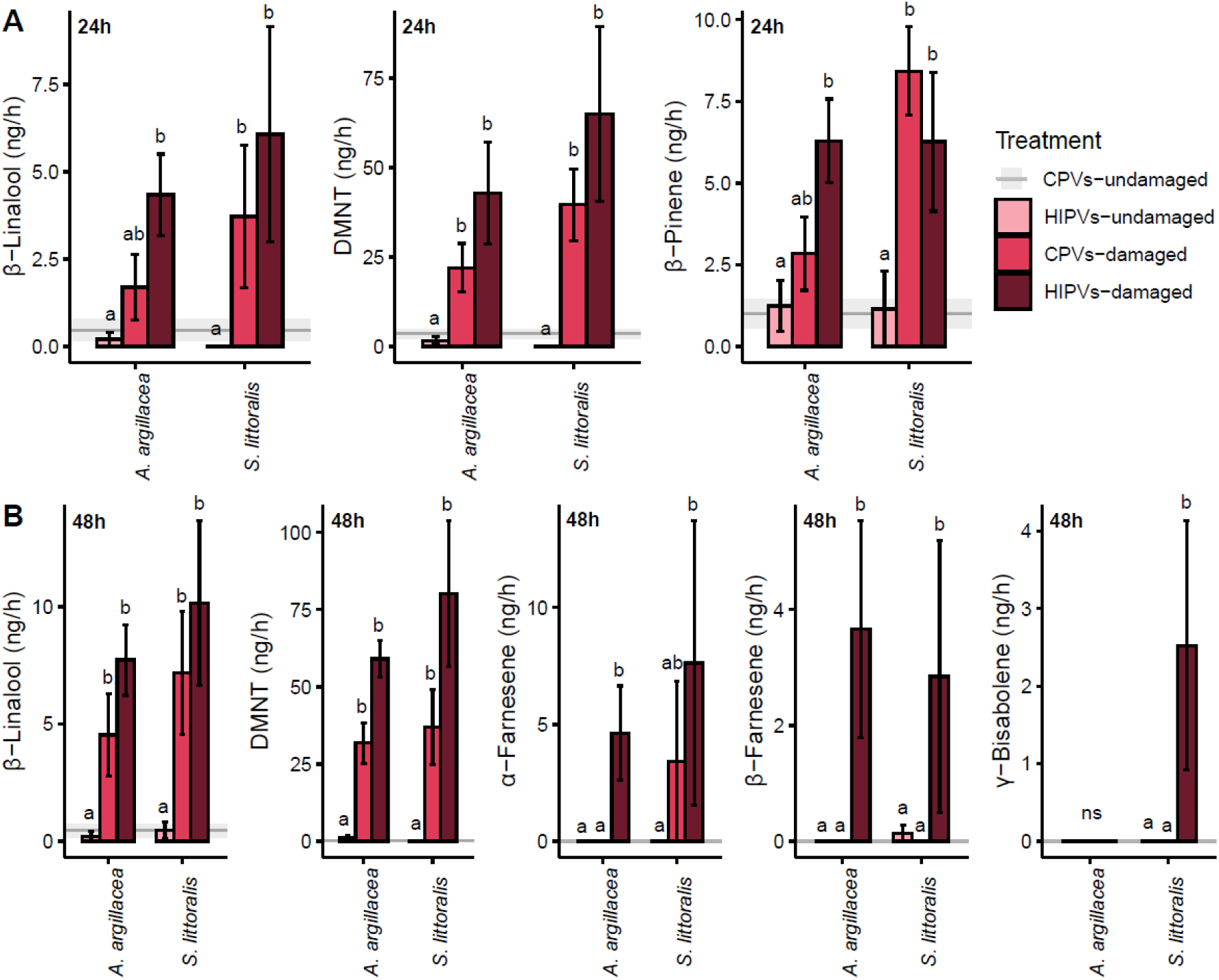
Mean volatile emissions (ng/hr ± SE) of individual compounds of differently treated receiver plants after **(A)** 24 h and **(B)** 48 h (n = 12). Receiver plants in 3^rd^ leaf stage were first exposed for 72 h to herbivore-induced plant volatiles from damaged emitter plants *(HIPVs*) or constitutive plant volatiles from undamaged emitter plants (*CPVs*), and were subsequently subjected to herbivory themselves (*damaged*) on the 1^st^ leaf for 48 h or left undamaged (*undamaged*). Different letters indicate significant differences (Tukey-adjusted pairwise comparisons). No differences were found between *HIPVs-undamaged* and *CPVs-undamaged* plants (baseline).

Phytohormones, gene expression and terpene aldehyde levels of receiver plants were analysed with the same approach as the volatile emission measurements. To test for direct induction, only the undamaged receiver plants were compared. To test for priming effects, the effect of treatment (HIPV exposure and/or insect damage) and the interaction with subsequent herbivory by the two insect species was tested.

Jasmonic acid (JA) and jasmonic acid-isoleucine (JA-Ile) levels were higher in undamaged plants exposed to HIPVs than in undamaged plants exposed to CPVs for both insects (JA: χ^2^_(2)_ = 54.095, p < 0.0001; JA-Ile: χ^2^_(2)_ = 27.789, p < 0.0001), with no differences between exposure to volatiles induced by *A. argillacea* or *S. littoralis* (Figure 4, Supplementary Table S5). Further, there was an overall effect of HIPVs exposure and/or damage treatment on JA-Ile levels, with no significant effect of insect species or the interaction and no overall effects for JA (JA: Treatment χ^2^_(2)_ = 2.7681, p = 0.08, Insect χ^2^_(1)_ = 3.0506, p = 0.09, Treatment:Insect χ^2^_(2)_ = 1.0964, p = 0.3; JA-Ile: Treatment χ^2^_(2)_ = 10.1506, p = 0.0062, Insect χ^2^_(1)_ = 2.1754, p = 0.14, Treatment:Insect χ^2^_(2)_ = 3.2295, p = 0.20). Undamaged receivers exposed to HIPVs induced by *A. argillacea* did not differ in JA or JA-Ile levels compared to damaged receivers, but for *S. littoralis*, damaged receivers that had first been exposed to HIPVs showed higher JA levels than damaged receivers exposed to CPVs, and higher JA-Ile levels than undamaged receivers exposed to HIPVs (Figure 4, Supplementary Table S5). For the other tested phytohormones (ABA, IAA, OPDA, SA), no significant differences were observed between any of the treatments (Supplementary Figure S3, Supplementary Table S6).

**Figure 4.**
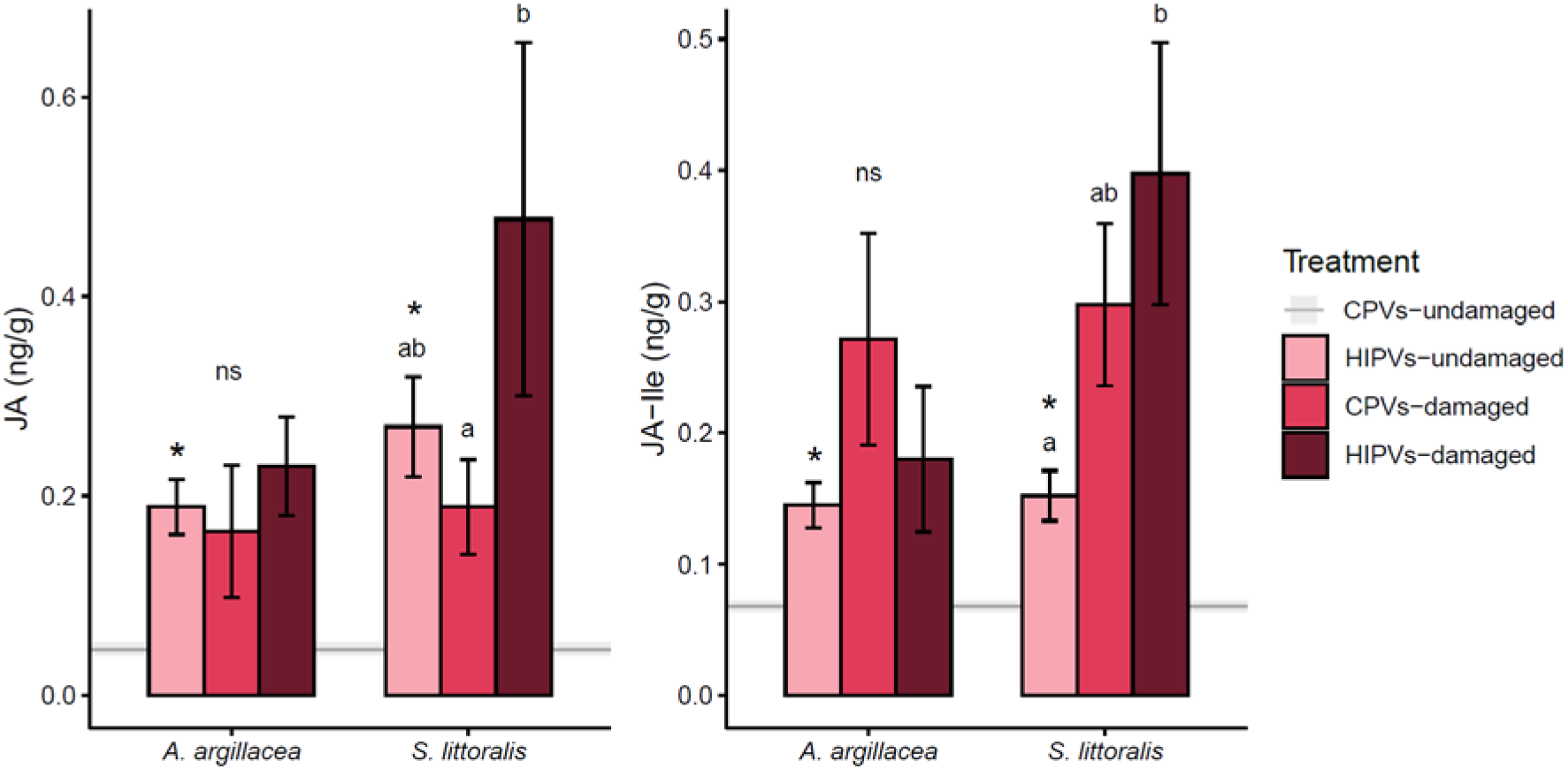
Mean levels (± SE) of phytohormones jasmonic acid (JA) and jasmonic acid-isoleucine (JA-Ile) in receiver plants (n = 5-7). Receiver plants in 3^rd^ leaf stage were first exposed for 72 h to herbivore-induced plant volatiles from damaged emitter plants *(HIPVs*) or constitutive plant volatiles from undamaged emitter plants (*CPVs*), and subsequently subjected to herbivory themselves (*damaged*) on the 1^st^ leaf for 24 h or left undamaged (*undamaged*). Phytohormones were then measured in the 3^rd^ leaf. Different letters indicate significant differences (Tukey-adjusted pairwise comparisons). Asterisks on the *HIPVs-undamaged* treatment indicate a significant difference compared to the *CPVs-undamaged* treatment (baseline).

In undamaged receiver plants, genes associated with the synthesis of terpene aldehydes were not differentially expressed comparing plants exposed to CPVs or HIPVs for both *2-ODD* and *Cdn1c3* (*2-ODD*: χ^2^_(2)_ = 0.5317, p = 0.77; *Cdn1c3*: χ^2^_(2)_ = 2.4952, p = 0.29, Figure 5A, Supplementary Table S7). In plants exposed to HIPVs and/or herbivory, there was an overall effect of treatment on gene expression but no difference between herbivory by the two insect species or the interaction (*2-ODD*: Treatment χ^2^_(2)_ = 34.628, p < 0.0001, Insect χ^2^_(1)_ = 1.454, p = 0.23, Treatment:Insect χ^2^_(2)_ = 3.2295, p = 0.89; *Cdn1c3*: Treatment χ^2^_(2)_ = 13.219, p = 0.0013, Insect χ^2^_(1)_ = 0.9378, p = 0.33, Treatment:Insect χ^2^_(2)_ = 0.0685, p = 0.97). Gene expression was significantly higher in damaged receivers than in undamaged receivers exposed to HIPVs, whereas in damaged receivers exposed to HIPVs gene expression was similar to that of the damaged receivers that had been exposed to CPVs (Figure 5A, Supplementary Table S7).

**Figure 5.**
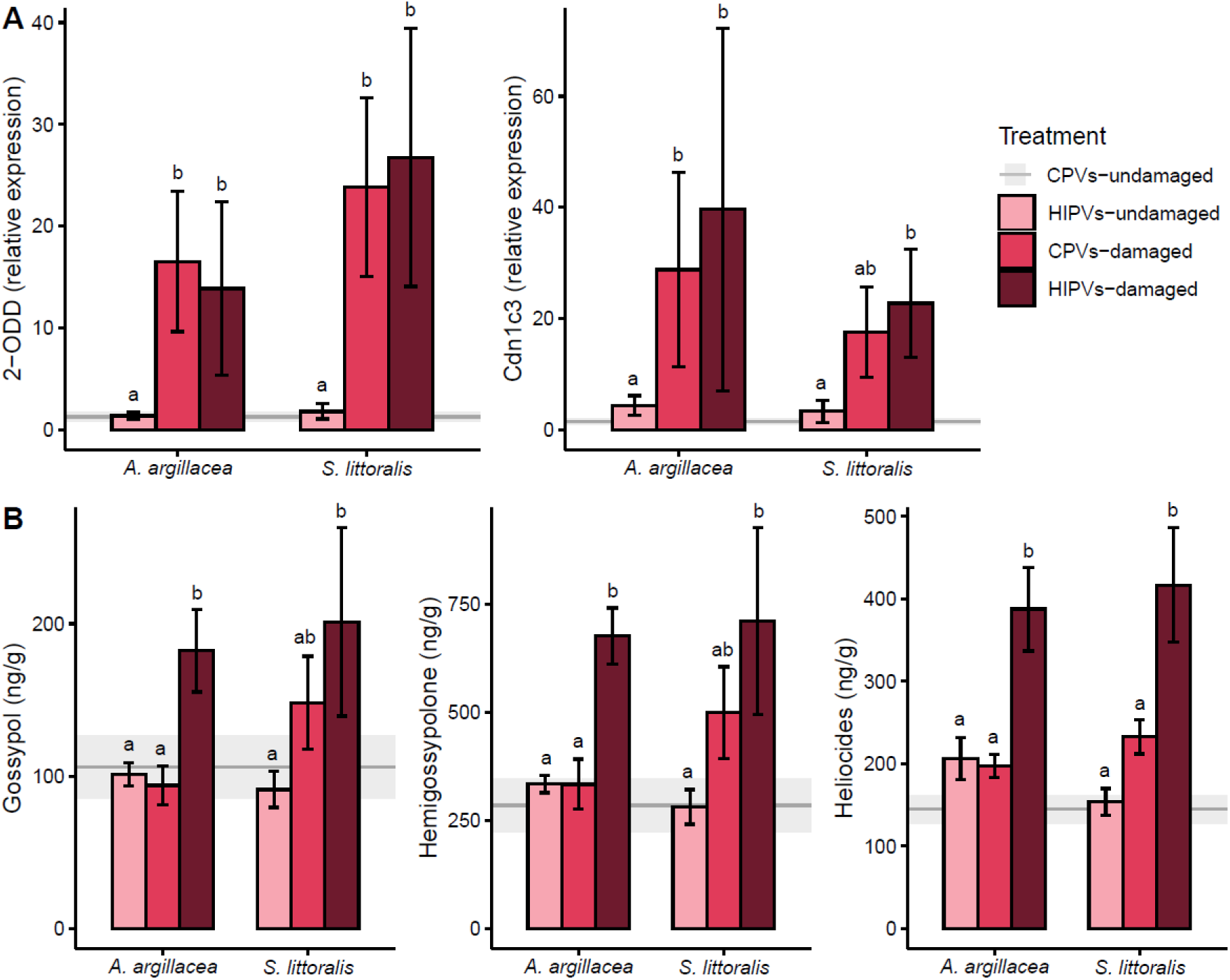
Mean levels (± SE) of **(A)** relative gene expression (n = 6 -12) and **(B)** terpene aldehydes in ng per g of fresh leaf mass (4^th^ leaf; n = 5-7) in receiver plants. Receiver plants in 3^rd^ leaf stage were first exposed for 72 h to herbivore-induced plant volatiles from damaged emitter plants *(HIPVs*) or constitutive plant volatiles from undamaged emitter plants (*CPVs*), and subsequently subjected to herbivory themselves (*damaged*) on the 1^st^ leaf or left undamaged (*undamaged*). After 24 h of receiver damage, gene expression was directly measured on the 3^rd^ leaf. For terpene aldehydes, receiver plants were damaged for 48 h, and then left for two weeks to develop the 4^th^ leaf. Different letters indicate significant differences (Tukey-adjusted pairwise comparisons). No differences were found between *HIPVs-undamaged* and *CPVs-undamaged* plants (baseline).

Accumulation of terpene aldehydes in leaves did not differ between undamaged plants exposed to HIPVs or CPVs (Gossypol: χ^2^_(2)_ = 0.5897, p = 0.74; Hemigossypolone: χ^2^_(2)_ = 0.78253, p = 0.68; Heliocides: χ^2^_(2)_ = 5.2995, p = 0.071, Figure 5B, Supplementary Table S8). In plants exposed to HIPVs and/or herbivory damage, there was an overall effect of treatment on levels of gossypol, hemigossypolone and heliocides, but not of insect species or the interaction (Gossypol: Treatment χ^2^_(2)_ = 26.915, p < 0.0001, Insect χ^2^_(1)_ = 0.951, p = 0.33, Treatment:Insect χ^2^_(2)_ = 5.592, p = 0.063; Hemigossypolone: Treatment χ^2^_(2)_ = 28.421, p < 0.0001, Insect χ^2^_(1)_ = 0.2616, p = 0.61, Treatment:Insect χ^2^_(2)_ = 3.6124, p = 0.16; Heliocides: Treatment χ^2^_(2)_ = 59.249, p < 0.0001, Insect χ^2^_(1)_ = 0.068, p = 0.79, Treatment:Insect χ^2^_(2)_ = 4.531, p = 0.1). Damaged receivers exposed to CPVs did not show different levels of terpene aldehydes than the undamaged receivers, but damaged receivers exposed to HIPVs had higher levels of gossypol and hemigossypolone for *A. argillacea*, and higher levels of heliocides for *A. argillacea* and *S. littoralis* compared to damaged receivers exposed to CPVs (Figure 5B, Supplementary Table S8).

When linking the matrix of volatiles of emitter plants to the terpene aldehyde levels in the receiver plants using a sPLS model, green leaf volatiles such as Z-3-hexenyl-2-methylbutyrate, Z-3-hexenyl acetate or Z-3-hexenol showed the strongest association with levels of gossypol, hemigossypolone and heliocides in receiver plants (R^2^ = 0.35, Figure 6A). For jasmonates, sesquiterpenes such as α-farnesene and nerolidol in the volatile blend of the emitter plants exhibited the strongest association with JA levels of receiver plants (R^2^ = 0.56, Figure 6B). JA-Ile was not strongly associated with volatiles of emitter plants (R^2^ = 0.10).

**Figure 6.**
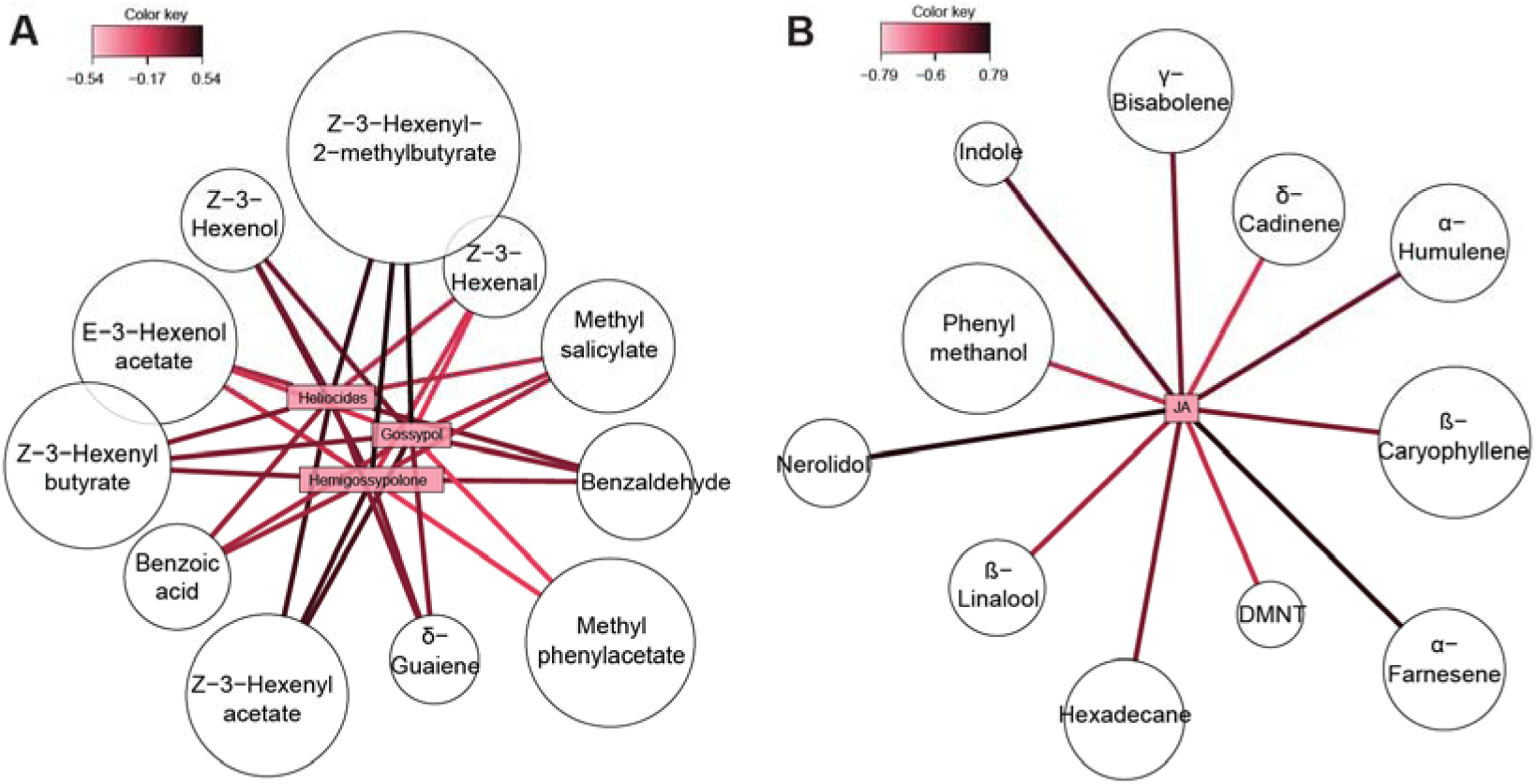
Network plots from sPLS models associating plant responses of receiver plants with the matrix of volatiles of emitter plants. **(A)** Network plot linking the volatile emissions of emitter plants to the levels of gossypol, hemigossypolone and heliocides in receiver plants (n = 5-7). **(B)** Network plot linking the volatile emissions of emitter plants to the JA level in receiver plants (n = 5-7). Colours represent the estimated correlation coefficient, and circle size is proportional to the importance of each volatile compound in the model (absolute loading value).

## Discussion

In this study, we compared the effects of herbivory on cotton plants by a monophagous and a highly polyphagous lepidopteran species on HIPV emissions and on plant-plant signalling resulting in defence priming. While priming effects in cotton have already been suggested for extrafloral nectar production (Briones-May *et al*., 2023; Quijano-Medina *et al*., 2024), we demonstrate here that priming effects occur for to the production of toxic and volatile defence metabolites, with distinct differences between the effects of HIPVs induced by a monophagous and a highly polyphagous caterpillar.

As expected, cotton plants attacked by *A. argillacea* or *S. littoralis* emitted more volatile compounds than undamaged control plants (Figure 1A). We had hypothesized that HIPV total emissions triggered by the monophagous herbivore *A. argillacea* might be less, as the suppression of HIPVs has been observed for specialist caterpillars in other plants (Sobhy *et al*., 2017; De Lange *et al*., 2020), but opposite effects are also known for some types of herbivores (Rowen & Kaplan, 2016). Although the total quantity of HIPVs after 72 h of attack did not differ between *A. argillacea* and *S. littoralis* (Figure 1A), there were differences in blend composition, with plants attacked by *A. argillacea* producing more benzoic acid and methyl salicylate, and plants attacked by *S. littoralis* emitting higher quantities of γ-bisabolene, caryophyllene oxide and decyne (Figure 1C). These differences might explain the observed differences in induction or priming effects in receiver plants.

When undamaged receiver plants were exposed to the HIPVs of emitter plants attacked by *A. argillacea* or *S. littoralis*, in both cases we observed an increase in the levels of jasmonates (Figure 4), but no direct induction of volatiles (Figure 2A), terpene aldehydes (Figure 5B) or expression of genes related to the terpene aldehydes (Figure 5A). Grandi et al. (2024) had observed an induction by HIPVs that resulted in increases in volatiles, phytohormones and expression of defensive genes in cotton plants immediately after HIPV exposure, but it remained unclear if this volatile-mediated induction of defences was relatively minor compared to the activation of a primed state (Martinez-Medina *et al*., 2016). To make the distinction between a small temporary defence increase or full-blown defence induction will greatly depend on timing of sample collection. In our study, phytohormone levels and gene expression were measured 24 h after the end of exposure to emitter volatiles, and terpene aldehydes levels were determined two weeks later. This timing of measurements was chosen to specifically distinguish between priming and induction, as they were performed when we expected to detect induction in response to herbivore damage. However, it is possible that direct changes in volatile emissions, gene expression or terpene aldehydes in response to HIPVs are transient and not detectable 24 h after volatile exposure. We found, however, comparable increases in jasmonates in plants exposed to HIPV only and plants subjected to insect damage only. When plants were exposed to both HIPVs and herbivory, the levels were still similar in the case of *A. argillacea* attack but significantly higher for *S. littoralis* attack (Figure 4). Either way, there was no evidence for priming of phytohormones since levels after HIPV-exposure and subsequent herbivory were not higher than expected from the addition of induced responses by herbivory and HIPVs separately. The results suggest that the levels of phytohormone induction to either HIPV exposure or mild insect damage was not enough to result in measurable induction of terpene aldehydes (Figure 5B); yet the increase in jasmonates caused by HIPV-exposure combined with herbivory clearly boosted the biochemical cascade involved in plant defence priming.

Hence, our results imply that exposure to HIPVs triggers a primed state in undamaged receiver plants, which is readily observable after they are damaged themselves. This was apparent from a strong defence induction than damaged receiver plants that had been exposed to CPVs of undamaged emitter plants. For receiver volatile emissions, the priming effect, although not statistically significant, resulted in apparent higher emissions of β-linalool, DMNT and β-pinene after 24 h of herbivory (Figure 3A), and after 48 h even more clearly of β-linalool and DMNT (Figure 3B). The priming effect was most evident for several sesquiterpenes, in the case of *S. littoralis* treatment only damaged primed plants emitted β-farnesene and γ-bisabolene, and in the *A. argillacea* treatment only damaged primed plants emitted α-farnesene and β-farnesene (Figure 3B). The induction of β-farnesene by primed plants in response to attacks by either herbivore species seems particularly relevant; it is known to be the alarm pheromone of aphids and act as a strong repellent against aphids when emitted by plants (Bowers *et al*., 1972; Wang *et al*., 2025). β-Farnesene has previously been reported to be released by plants after herbivory by lepidopteran species (Turlings *et al*., 1990; Loughrin *et al*., 1994) and has potential insecticidal effects (Sun *et al*., 2022). Volatile emissions of cotton plants in a primed state may also influence the oviposition and feeding behaviour of herbivorous insects. For instance, *Spodoptera littoralis* females lay more eggs on undamaged cotton plants exposed to volatiles from undamaged neighbour plants than on plants exposed to HIPVs (Zakir *et al*., 2013), and *S. exigua* caterpillars much prefer leaves from cotton plants exposed to volatiles from undamaged neighbouring plants than to those exposed to volatiles from damaged neighbours (Grandi *et al*., 2024). Taken together, these studies show that herbivores can recognize and avoid plants in a primed state through their volatile profile, thereby circumventing exposure to highly induced defences in these plants, which in turn protects these HIPV-primed plants. Thus, exposure to volatiles alone might be as efficient as induction by herbivory regarding plant protection (Llandres *et al*., 2018).

The differences found in plant responses to the two herbivore species suggests some degree of herbivore specificity. The priming of γ-bisabolene emissions and the higher induction of JA and JA-Ile in primed receiver plants in the *S. littoralis* treatment could be linked to a faster induction of HIPVs and a stronger induction in response to *S. littoralis* damage or, alternatively, to a suppressive effect of volatiles specifically induced by *A. argillacea*. Although this might have implications for subsequent interactions with each herbivore species, the final defensive response (levels of terpene aldehydes in primed plants) was found to be similar for both herbivore species. Thus, it seems that in this case the actual priming of chemical defences is a general response to herbivory that is independent of the herbivore identity and diet breath.

Induction by damage after priming resulted in terpene aldehydes levels up to 100% higher than in non-primed plants (Figure 5B). However, we found no difference in terpenoid aldehydes between undamaged receivers exposed to HIPVs and undamaged receivers exposed to CPVs, i.e., no evidence of direct induction. This is in contrast with the direct induction by HIPVs reported previously of ∼20% (Grandi *et al*., 2024), and suggests that in our case exposure to HIPVs alone was not enough to increase the concentration of terpene aldehydes after two weeks, which is the reported time that it takes for new leaves to accumulate these compounds (Bezemer *et al*., 2004). Herbivory without prior HIPV exposure also did not increase terpene aldehyde concentrations, possibly because a single caterpillar only causes minor damage in 24 h. This indicates that plants only respond to fresh and mild damage when they are primed by HIPVs from nearby attacked plants. This is particularly relevant in an ecological context since terpene aldehydes can have toxic, antimicrobial, antiviral and insecticidal effects (Stipanovic *et al*., 1988; Tian *et al*., 2016) and can also act as feeding deterrents (Meisner *et al*., 1977) thus potentially providing protection against a wide range of antagonists.

It is known that exposure to volatiles of wild cotton attacked by *S. frugiperda* augments the induction in response to damage of extra-floral nectar (EFN) (Briones-May *et al*., 2023) and heliocides (Quijano-Medina *et al*., 2024). A similar increase in the induction in EFN concentration after damage was observed in wild cotton plants exposed to volatiles of plants attacked by *A. argillacea* when emitter herbivory levels were high, i.e., 3 (vs 1) caterpillars per plant (Interian-Aguiñaga *et al*., 2025). Within this context, the present study seems to be the first to show HIPV-mediated priming effects in cotton for different defence pathways (volatile emissions and terpene aldehydes) activated via jasmonate induction. The present study and the one by (Grandi *et al*., 2024) demonstrate that HIPVs can trigger a minor short-term increase in terpene aldehydes and a substantial increase in phytohormones of the jasmonic pathway, this in turn results in the priming of several key plant volatiles and terpene aldehydes, but only after herbivore damage. This is in line with the simultaneous induction and priming of plant volatiles upon exposure to HIPVs in maize plants (Waterman *et al*., 2024) but extended to defence compounds that accumulate in the leaves.

Whether the entire volatile blend of the emitter plant or only specific compounds cause the priming effect on the receiver plants remains clear. Grandi et al. (2024) showed that defence induction in cotton by HIPVs is mediated by *de novo* synthesised volatiles (Grandi *et al*., 2024). Linking emitter HIPVs to the levels of terpene aldehydes and JA in receiver plants showed that terpene aldehyde induction seemed mainly linked to GLVs such as Z-3-hexenyl acetate and Z-3-hexenyl-2-methyl butyrate, whereas JA levels were associated with sesquiterpenes such as α-farnesene and nerolidol (Figure 6). Indeed, Z-3-hexenyl acetate, Z-3-hexenyl-2-methyl butyrate and α-farnesene are inducible and *de novo* synthesised after herbivore attack and they are released systemically from attacked plants (Röse *et al*., 1996). GLVs also have been shown to prime for higher JA levels and volatile release in maize (Engelberth *et al*., 2004), and to induce EFN production in lima bean (Heil *et al*., 2008). Other studies found GLVs as well as monoterpenes, sesquiterpenes and hemiterpenes to prime defences in maize, lima bean, sagebrush, and tea (Arimura *et al*., 2002; Kessler *et al*., 2006; Ton *et al*., 2007; Zhao *et al*., 2020; Midzi *et al*., 2022). Among studies testing for volatile mixture vs. individual compound effects, Kikuta et al. (2011) showed that *Chrysanthemum cinerariaefolium* plants induced pyrethrin levels when exposed to the volatile blend of damaged plants, but did not react when they were exposed to individual compounds of the same mixture (Kikuta *et al*., 2011). Indeed, volatile blends with specific ratios seem to play important roles in defence induction, and the importance of the combination of multiple compounds and their specificity is still an open question (Bouwmeester *et al*., 2019; Rosenkranz *et al*., 2021), especially in cotton plants.

With this study we show that exposure to HIPVs triggered by two herbivore species with contrasting diet breadth, *S. littoralis* and *A. argillacea*, prime multiple defence traits in cotton plants. These findings contribute to the overall understanding of plant-plant signalling and its consequences for antiherbivore defences. Further research with multiple specialist and generalist herbivores is still needed to determine if they induce specific blends of HIPVs that may have distinct priming effects on exposed plants and to gain more insight into which compounds are responsible for priming in cotton.

## Data availability

Data and R codes are available at Zenodo at DOI: https://doi.org/10.5281/zenodo.16570554.

## Author contributions

TCJT, CBS and LAR conceived the study. CBS, TCTJ and KA designed the experiments. KA, CBS, WY and AV performed the experiments and processed the data. LAR provided the *A. argillacea* insects. KA analysed the data and wrote the first manuscript. All authors revised the manuscript.

## Acknowledgements

The authors thank Juan Traine for his help with experimental manipulations, Marine Mamin and Mary V. Clancy for their help with the insect rearing, Thomas Degen for the experimental design illustration, and all members of the FARCE group at the University of Neuchâtel for helpful discussions.

