## Supporting information for "Exposure to herbivore-induced plant volatiles directly induces jasmonic acid and primes chemical defences in cotton plants"

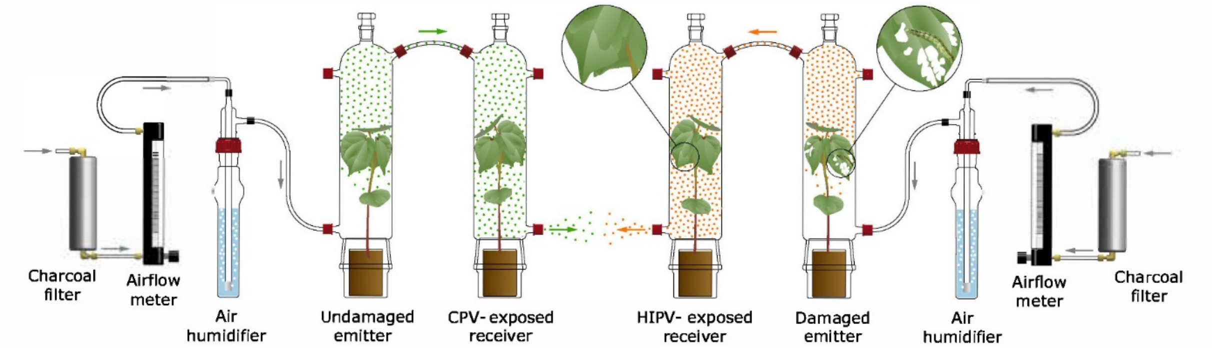


**Figure S1.** Experimental setup for the volatile exposure. Left side shows a receiver plant exposed to CPVs from an undamaged cotton plant, right side shows a receiver plant exposed to HIPVs from an *A. argillacea* - damaged cotton plant.

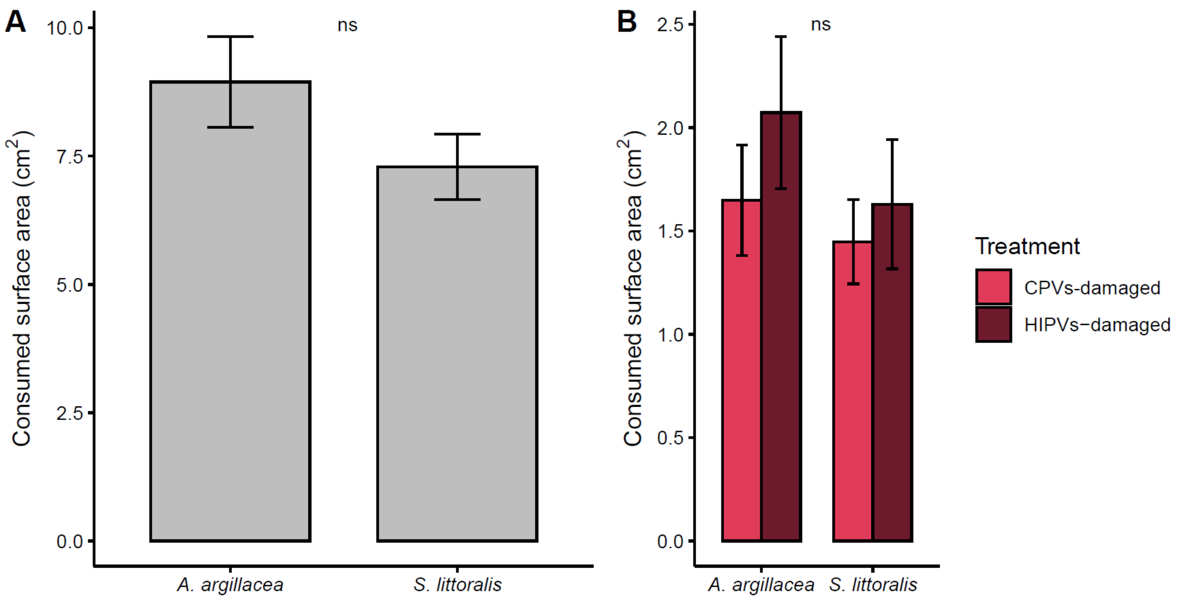


**Figure S2.** Mean consumed leaf surface area (± SE) of **(A)** emitter plants after 72 h of infestation by insects (n = 24) or **(B)** receiver plants after 48 h of infestation by insects (n = 6). NS indicates no significant difference among the treatments (Tukey-adjusted pairwise comparisons).

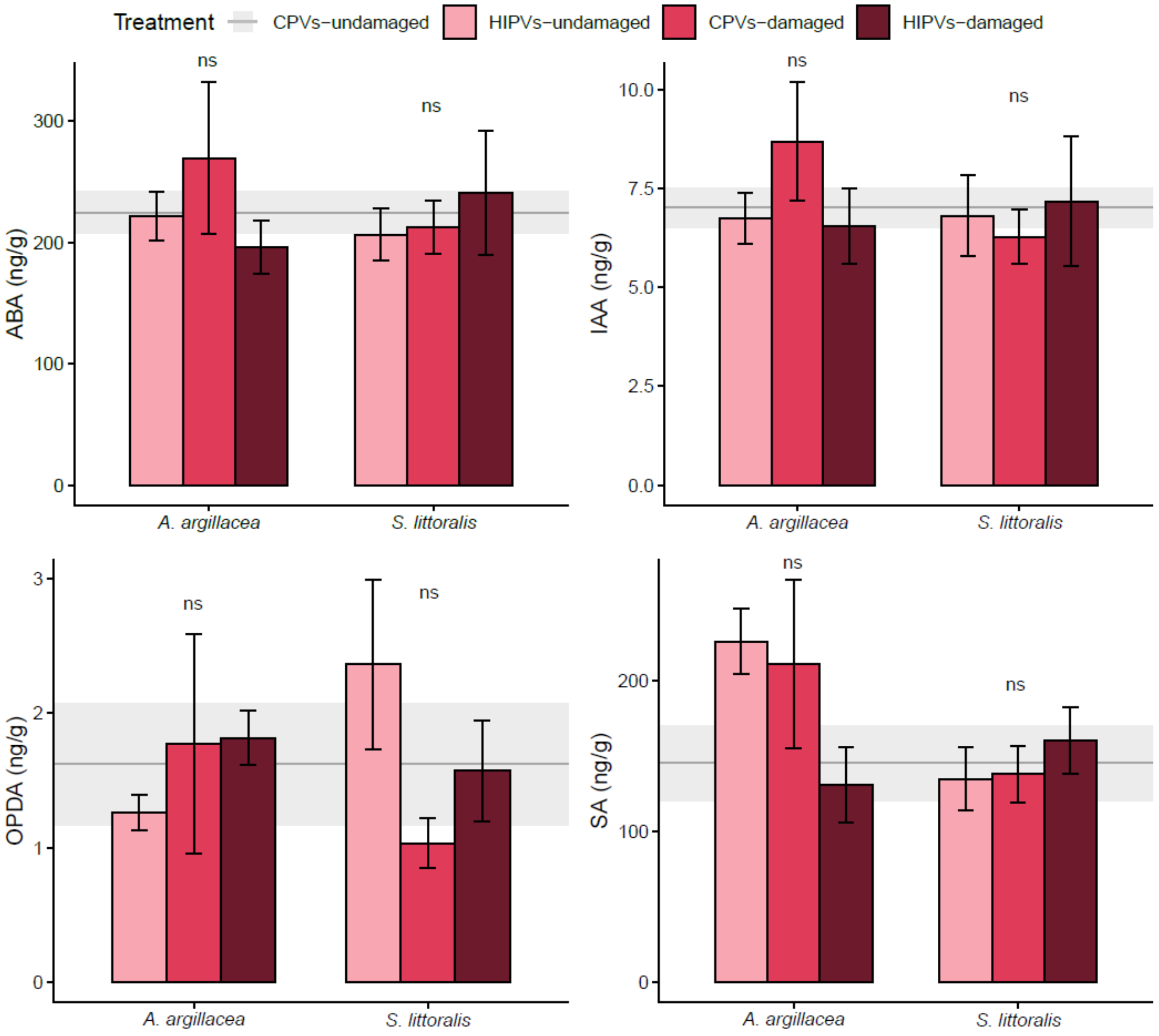


**Figure S3.** Mean levels (± SE) of the phytohormones abscisic acid (ABA), indole-3-acetic acid (IAA), 12-oxophytodienoic acid (OPDA) and salicylic acid (SA) in receiver plants (n = 5-7). Receiver plants in 3$^{rd}$ leaf stage were first exposed for 72 h to herbivore-induced plant volatiles from damaged emitter plants *(HIPVs)* or constitutive plant volatiles from undamaged emitter plants (*CPVs*), and subsequently subjected to herbivory themselves (*damaged*) on the 1$^{st}$ leaf for 24 h or left undamaged (*undamaged*). Phytohormones were then measured in the 3$^{rd}$ leaf. Different letters indicate significant differences (Tukey-adjusted pairwise comparisons). No differences were found between *HIPVs-undamaged* and *CPVs-undamaged* plants (baseline).

**Table S1.** Mean volatile emissions (± SE) of emitter plants after 72 h of infestation. P-values were adjusted (false discovery rate correction). Different letters indicate significant differences between treatments (Tukey-adjusted pairwise comparisons).

| **Compound** | **Undamaged** | ***A. argillacea*** | ***S. littoralis*** | **p-value** |
|---|---|---|---|---:|
| α-Copaene | 0 ± 0 a | 0.4 ± 0.17 b | 0.82 ± 0.25 b | < 0.0001 |
| α-Farnesene | 0 ± 0 a | 8.63 ± 1.69 b | 8.13 ± 3.15 b | < 0.0001 |
| α-Guaiene | 0 ± 0 a | 0.61 ± 0.61 a | 0.27 ± 0.2 a | 0.3 |
| α-Humulene | 0 ± 0 a | 1.26 ± 1.23 a | 0.62 ± 0.35 a | 0.2 |
| Alloaromadendrene | 0 ± 0 a | 0.3 ± 0.15 ab | 0.38 ± 0.2 b | 0.03 |
| α-Pinene | 1.62 ± 0.4 a | 66.84 ± 17.54 b | 85.39 ± 25.53 b | < 0.0001 |
| β-Caryophyllene | 0 ± 0 a | 5.04 ± 4.34 b | 5.54 ± 2.05 b | 0.002 |
| Benzaldehyde | 4.08 ± 0.77 a | 4.9 ± 0.57 a | 5.07 ± 0.67 a | 0.6 |
| Benzoic acid | 0 ± 0 a | 4.16 ± 0.86 b | 1.84 ± 0.63 c | < 0.0001 |
| Benzothiazole | 0.07 ± 0.03 a | 0.01 ± 0.01 a | 0.09 ± 0.05 a | 0.2 |
| β-Farnesene | 0.01 ± 0.01 a | 6.61 ± 0.95 b | 9.08 ± 2.26 b | < 0.0001 |
| β-Linalool | 0.16 ± 0.08 a | 39.26 ± 8.08 b | 45.9 ± 11.86 b | < 0.0001 |
| β-Myrcene | 0.02 ± 0.01 a | 16.44 ± 3.53 b | 10.07 ± 3.42 b | < 0.0001 |
| β-Ocimene | 0.82 ± 0.59 a | 173.79 ± 31.93 b | 118.87 ± 31.08 b | < 0.0001 |
| β-Phenylethyl acetate | 0 ± 0 a | 1.5 ± 0.31 b | 1.49 ± 0.37 b | < 0.0001 |
| β-Pinene | 0.23 ± 0.08 a | 7.95 ± 2.24 b | 10.05 ± 3.38 b | < 0.0001 |
| Caryophyllene oxide | 0.68 ± 0.29 a | 5.81 ± 1.81 b | 14.96 ± 4.54 c | < 0.0001 |
| δ-Cadinene | 0 ± 0 a | 0.85 ± 0.27 b | 1.21 ± 0.39 b | < 0.0001 |
| 1-Decyne | 4.26 ± 0.62 a | 8.47 ± 1.82 a | 35.97 ± 13.77 b | < 0.0001 |
| δ-Guaiene | 0 ± 0 a | 1.42 ± 1.41 a | 0.37 ± 0.27 a | 0.1 |
| DMNT | 5.96 ± 1.38 a | 86.3 ± 14.19 b | 64.59 ± 13.73 b | < 0.0001 |
| Hexadecane | 0 ± 0 a | 0.1 ± 0.09 a | 0.02 ± 0.02 a | 0.3 |
| E-2-Hexenal | 0.67 ± 0.33 a | 24.18 ± 10.82 b | 51.63 ± 16.79 b | < 0.0001 |
| Z-3-Hexenal | 1.47 ± 0.74 a | 37.71 ± 11.55 b | 69.79 ± 18.19 b | < 0.0001 |
| Z-3-Hexenol | 0.2 ± 0.09 a | 15.65 ± 4.72 b | 18.26 ± 7.19 b | < 0.0001 |
| E-3-Hexenol acetate | 0.09 ± 0.05 a | 2.77 ± 0.8 b | 2.86 ± 1.02 b | < 0.0001 |
| E-2-Hexenyl-2-methylbutyrate | 0.06 ± 0.06 a | 1.66 ± 0.47 b | 0.96 ± 0.23 b | < 0.0001 |
| Z-3-Hexenyl acetate | 4.92 ± 1.96 a | 61.86 ± 16.02 b | 48.68 ± 14.83 b | < 0.0001 |
| Z-3-Hexenyl butyrate | 0.03 ± 0.02 a | 1.95 ± 0.71 b | 1.03 ± 0.37 b | < 0.0001 |
| Z-3-Hexenyl-2-methylbutyrate | 0 ± 0 a | 0.97 ± 0.22 b | 0.56 ± 0.17 b | < 0.0001 |
| Indole | 0.13 ± 0.1 a | 79.12 ± 16.69 b | 98.72 ± 29.69 b | < 0.0001 |
| Jasmone | 0 ± 0 a | 4.66 ± 0.89 b | 3.69 ± 1.25 b | < 0.0001 |
| Limonene | 0.13 ± 0.05 a | 3.87 ± 1.16 b | 5.53 ± 1.66 b | < 0.0001 |
| Methyl anthranilate | 0 ± 0 a | 0.4 ± 0.11 b | 0.2 ± 0.08 b | < 0.0001 |
| Methyl phenylacetate | 1.38 ± 0.11 a | 1.66 ± 0.12 ab | 2.04 ± 0.32 b | 0.04 |
| Methyl salicylate | 0.52 ± 0.23 ab | 1.36 ± 0.41 a | 0.34 ± 0.14 b | 0.03 |
| Myroxide | 0 ± 0 a | 7.34 ± 1.69 b | 4.66 ± 1 b | < 0.0001 |
| Nerolidol | 0.01 ± 0.01 a | 2.08 ± 0.42 b | 2.24 ± 1.07 b | < 0.0001 |
| Nonanal | 8.31 ± 1.99 a | 8.04 ± 1.96 a | 8.31 ± 1.52 a | 0.9 |
| Pentadecane | 0 ± 0 a | 0.01 ± 0.01 a | 1.87 ± 1.15 b | 0.0001 |
| Phenylmethanol | 0.01 ± 0.01 a | 0.49 ± 0.18 a | 1.45 ± 0.57 b | < 0.0001 |
| TMTT | 5.64 ± 2 a | 9.93 ± 2.26 a | 6.81 ± 2.18 a | 0.4 |
| γ-Bisabolene | 0 ± 0 a | 0.25 ± 0.2 a | 1.99 ± 1 b | < 0.0001 |
| γ-Elemene | 0 ± 0 a | 0.19 ± 0.16 ab | 0.68 ± 0.33 b | 0.008 |

**Table S2.** Complete statistics for the total volatile emissions of receiver plants, supplementing Figure 3A. Direct induction was tested by comparing only the undamaged receiver plants exposed to CPVs or HIPVs from *A. argillacea* or *S. littoralis*. Priming effects were tested by testing for effects of treatment (HIPV exposure and /or insect damage) and the interaction with the insect species. Post-hoc tests were performed as tukey-adjusted pairwise comparisons.

| Volatile emissions | Test | Comparison | Test value | Df | p-value |
|---|---|---|---|---|---|
| **Testing for induction** | | | | | |
| Total emissions 24 h | Anova | Treatment | $\chi^2 = 0.0109$ | 1 | 0.91 |
| Total emissions 48 h | Anova | Treatment | $\chi^2 = 0.2596$ | 1 | 0.61 |
| **Testing for priming effects** | | | | | |
| Total emissions 24 h | Anova | Treatment | $\chi^2 = 12.435$ | 2 | 0.002 |
| | | Insect | $\chi^2 = 0.0172$ | 1 | 0.9 |
| | | Treatment:Insect | $\chi^2 = 4.1761$ | 2 | 0.1 |
| | Post-hoc | HIPVs-undamaged vs. CPVs-undamaged (*A. argillacea*) | t = -0.726 | 30 | 0.75 |
| | | HIPVs-undamaged vs. HIPVs-damaged (*A. argillacea*) | t = -1.500 | 30 | 0.31 |
| | | CPVs-damaged vs. HIPVs-damaged (*A. argillacea*) | t = -0.774 | 30 | 0.72 |
| | | HIPVs-undamaged vs. CPVs-undamaged (*S. littoralis*) | t = -3.526 | 30 | 0.0038 |
| | | HIPVs-undamaged vs. HIPVs-damaged (*S. littoralis*) | t = -3.558 | 30 | 0.0035 |
| | | CPVs-damaged vs. HIPVs-damaged (*S. littoralis*) | t = -0.032 | 30 | 0.99 |
| Total emissions 48 h | Anova | Treatment | $\chi^2 = 67.199$ | 2 | < 0.0001 |
| | | Insect | $\chi^2 = 0.121$ | 1 | 0.73 |
| | | Treatment:Insect | $\chi^2 = 2.356$ | 2 | 0.31 |
| | Post-hoc | HIPVs-undamaged vs. CPVs-undamaged (*A. argillacea*) | t = -4.449 | 30 | 0.0003 |
| | | HIPVs-undamaged vs. HIPVs-damaged (*A. argillacea*) | t = -4.730 | 30 | 0.0001 |
| | | CPVs-damaged vs. HIPVs-damaged (*A. argillacea*) | t = -0.281 | 30 | 0.96 |
| | | HIPVs-undamaged vs. CPVs-undamaged (*S. littoralis*) | t = -5.553 | 30 | < 0.0001 |
| | | HIPVs-undamaged vs. HIPVs-damaged (*S. littoralis*) | t = -6.902 | 30 | < 0.0001 |
| | | CPVs-damaged vs. HIPVs-damaged (*S. littoralis*) | t = -1.350 | 30 | 0.38 |

**Table S3.** Pairwise RDA tests (999 permutations) with adjusted p-values (Holm method) supplementing Figure 3B.

| **Comparison** | **F-value** | **Df** | **p-value** |
|---|---|---|---|
| ***A. argillacea* 24 h** | | | |
| CPVs-undamaged vs. HIPVs-undamaged | 0.6408 | 1 | 1.000 |
| CPVs-undamaged vs. CPVs-damaged | 2.042 | 1 | 0.240 |
| CPVs-undamaged vs. HIPVs-damaged | 4.0000 | 1 | 0.024 |
| HIPVs-undamaged vs. CPVs-damaged | 1.3068 | 1 | 0.726 |
| HIPVs-undamaged v.s HIPVs-damaged | 2.4686 | 1 | 0.145 |
| CPVs-damaged vs. HIPVs-damaged | 0.794 | 1 | 1.000 |
| ***S. littoralis* 24 h** | | | |
| CPVs-undamaged vs. HIPVs-undamaged | 0.9521 | 1 | 0.880 |
| CPVs-undamaged vs. CPVs-damaged | 3.357 | 1 | 0.024 |
| CPVs-undamaged vs. HIPVs-damaged | 2.9338 | 1 | 0.100 |
| HIPVs-undamaged vs. CPVs-damaged | 3.4375 | 1 | 0.025 |
| HIPVs-undamaged vs. HIPVs-damaged | 2.5583 | 1 | 0.267 |
| CPVs-damaged vs. HIPVs-damaged | 0.5039 | 1 | 0.885 |
| ***A. argillacea* 48 h** | | | |
| CPVs-undamaged vs. HIPVs-undamaged | 0.6601 | 1 | 1.000 |
| CPVs-undamaged vs. CPVs-damaged | 4.3387 | 1 | 0.025 |
| CPVs-undamaged vs. HIPVs-damaged | 5.1322 | 1 | 0.025 |
| HIPVs-undamaged vs. CPVs-damaged | 3.3002 | 1 | 0.039 |
| HIPVs-undamaged vs. HIPVs-damaged | 3.7973 | 1 | 0.024 |
| CPVs-damaged vs. HIPVs-damaged | 0.4393 | 1 | 1.000 |
| ***S. littoralis* 48 h** | | | |
| CPVs-undamaged vs. HIPVs-undamaged | 0.5094 | 1 | 1.000 |
| CPVs-undamaged vs. CPVs-damaged | 2.9491 | 1 | 0.032 |
| CPVs-undamaged vs. HIPVs-damaged | 3.9178 | 1 | 0.012 |
| HIPVs-undamaged vs. CPVs-damaged | 2.335 | 1 | 0.036 |
| HIPVs-undamaged vs. HIPVs-damaged | 3.0657 | 1 | 0.020 |
| CPVs-damaged vs. HIPVs-damaged | 0.7968 | 1 | 1.000 |

**Table S4.** Complete statistics for the individual compounds of receiver volatile emissions shown in Figure 4. Direct induction was tested by comparing only the undamaged receiver plants exposed to CPVs or HIPVs from *A. argillacea* or *S. littoralis*. Priming effects were tested by testing for effects of treatment (HIPV exposure and /or insect damage) and the interaction with the insect species. Post-hoc tests were performed as tukey-adjusted pairwise comparisons.

| **Compound** | **Test** | **Comparison** | **Test value** | **Df** | **p-value** |
|---|---|---|---|---|---|
| **Testing for induction** | | | | | |
| β-Linalool 24 h | Anova | Treatment | $\chi^2$ = 2.16 | 1 | 0.14 |
| β-Linalool 48 h | Anova | Treatment | $\chi^2$ = 0.11324 | 1 | 0.74 |
| DMNT 24 h | Anova | Treatment | $\chi^2$ = 3.6015 | 1 | 0.058 |
| DMNT 48 h | Anova | Treatment | $\chi^2$ = 0.30688 | 1 | 0.58 |
| β-Pinene 24 h | Anova | Treatment | $\chi^2$ = 0.03892 | 1 | 0.84 |
| β-Farnesene | Anova | Treatment | $\chi^2$ = 0.51421 | 1 | 0.47 |
| **Testing for priming effect** | | | | | |
| β-Linalool 24 h | Anova | Treatment | $\chi^2$ = 27.0301 | 2 | < 0.0001 |
| | | Insect | $\chi^2$ = 0.7269 | 1 | 0.39 |
| | | Treatment:Insect | $\chi^2$ = 1.4831 | 2 | 0.48 |
| β-Linalool 48 h | Anova | Treatment | $\chi^2$ = 50.648 | 2 | < 0.0001 |
| | | Insect | $\chi^2$ = 1.623 | 1 | 0.20 |
| | | Treatment:Insect | $\chi^2$ = 0.154 | 2 | 0.93 |
| | Post-hoc | HIPVs-undamaged vs. CPVs-undamaged (*A. argillacea*) | t = -4.076 | 30 | 0.0009 |
| | | HIPVs-undamaged vs. HIPVs-damaged (*A. argillacea*) | t = -5.297 | 30 | < 0.0001 |
| | | CPVs-damaged vs. HIPVs-damaged (*A. argillacea*) | t = -1.222 | 30 | 0.45 |
| | | HIPVs-undamaged vs. CPVs-undamaged (*S. littoralis*) | t = -4.613 | 30 | 0.0002 |
| | | HIPVs-undamaged vs. HIPVs-damaged (*S. littoralis*) | t = -5.443 | 30 | < 0.0001 |
| | | CPVs-damaged vs. HIPVs-damaged (*S. littoralis*) | t = -0.830 | 30 | 0.69 |
| DMNT 24 h | Anova | Treatment | $\chi^2$ = 86.469 | 2 | < 0.0001 |
| | | Insect | $\chi^2$ = 0.004 | 1 | 0.95 |
| | | Treatment:Insect | $\chi^2$ = 7.619 | 2 | 0.022 |
| DMNT 48 h | Anova | Treatment | $\chi^2$ = 168.453 | 2 | < 0.0001 |
| | | Insect | $\chi^2$ = 0.179 | 1 | 0.67 |
| | | Treatment:Insect | $\chi^2$ = 5.091 | 2 | 0.78 |
| | Post-hoc | HIPVs-undamaged vs. CPVs-undamaged (*A. argillacea*) | t = -4.076 | 30 | < 0.0001 |
| | | HIPVs-undamaged vs. HIPVs-damaged (*A. argillacea*) | t = -5.297 | 30 | < 0.0001 |
| | | CPVs-damaged vs. HIPVs-damaged (*A. argillacea*) | t = -1.222 | 30 | 0.20 |
| | | HIPVs-undamaged vs. CPVs-undamaged (*S. littoralis*) | t = -4.613 | 30 | < 0.0001 |
| | | HIPVs-undamaged vs. HIPVs-damaged (*S. littoralis*) | t = -5.443 | 30 | < 0.0001 |
| | | CPVs-damaged vs. HIPVs-damaged (*S. littoralis*) | t = -0.830 | 30 | 0.086 |
| β-Pinene 24 h | Anova | Treatment | $\chi^2$ = 13.7870 | 2 | 0.001 |
| | | Insect | $\chi^2$ = 1.0559 | 1 | 0.30 |
| | | Treatment:Insect | $\chi^2$ = 2.6456 | 2 | 0.27 |
| α- Farnesene 48 h | Anova | Treatment | $\chi^2$ = 17.3952 | 2 | 0.00017 |
| | | Insect | $\chi^2$ = 2.6172 | 1 | 0.11 |
| | | Treatment:Insect | $\chi^2$ = 2.7994 | 2 | 0.25 |
| | Post-hoc | HIPVs-undamaged vs. CPVs-undamaged (*A. argillacea*) | t = 0.000 | 30 | 1.00 |
| | | HIPVs-undamaged vs. HIPVs-damaged (*A. argillacea*) | t = -2.703 | 30 | 0.029 |
| | | CPVs-damaged vs. HIPVs-damaged (*A. argillacea*) | t = -2.703 | 30 | 0.029 |
| | | HIPVs-undamaged vs. CPVs-undamaged (*S. littoralis*) | t = -2.326 | 30 | 0.068 |
| | | HIPVs-undamaged vs. HIPVs-damaged (*S. littoralis*) | t = -3.372 | 30 | 0.0057 |
| | | CPVs-damaged vs. HIPVs-damaged (*S. littoralis*) | t = -1.046 | 30 | 0.55 |
| β-Farnesene 48 h | Anova | Treatment | $\chi^2$ = 31.322 | 2 | < 0.0001 |
| | | Insect | $\chi^2$ = 0.0071 | 1 | 0.93 |
| | | Treatment:Insect | $\chi^2$ = 0.2848 | 2 | 0.87 |
| | | HIPVs-undamaged vs. CPVs-undamaged (*A. argillacea*) | t = 0.000 | 30 | 1.00 |

| | | | | | |
|---|---|---|---|---|---|
| | Post-hoc | HIPVs-undamaged vs. HIPVs-damaged (*A. argillacea*) | t = -3.594 | 30 | 0.0032 |
| | | CPVs-damaged vs. HIPVs-damaged (*A. argillacea*) | t = -3.594 | 30 | 0.0032 |
| | | HIPVs-undamaged vs. CPVs-undamaged (*S. littoralis*) | t = 0.302 | 30 | 0.95 |
| | | HIPVs-undamaged vs. HIPVs-damaged (*S. littoralis*) | t = -2.844 | 30 | 0.021 |
| | | CPVs-damaged vs. HIPVs-damaged (*S. littoralis*) | t = -3.146 | 30 | 0.010 |
| γ-Bisabolene 48 h | Anova | Treatment | $\chi^2$ = 18.0668 | 2 | 0.00012 |
| | | Insect | $\chi^2$ = 6.5872 | 1 | 0.010 |
| | | Treatment:Insect | $\chi^2$ = 14.9285 | 2 | 0.00057 |
| | Post-hoc | HIPVs-undamaged vs. CPVs-undamaged (*A. argillacea*) | t = 0.000 | 30 | 1.00 |
| | | HIPVs-undamaged vs. HIPVs-damaged (*A. argillacea*) | t = 0.000 | 30 | 1.00 |
| | | CPVs-damaged vs. HIPVs-damaged (*A. argillacea*) | t = 0.000 | 30 | 1.00 |
| | | HIPVs-undamaged vs. CPVs-undamaged (*S. littoralis*) | t = 0.000 | 30 | 1.00 |
| | | HIPVs-undamaged vs. HIPVs-damaged (*S. littoralis*) | t = -4.784 | 30 | 0.0001 |
| | | CPVs-damaged vs. HIPVs-damaged (*S. littoralis*) | t = -4.784 | 30 | 0.0001 |

**Table S5.** Complete statistics for the phytohormones jasmonic acid (JA) and jasmonic acid-isoleucine (JA-Ile) shown in Figure 5. Direct induction was tested by comparing only the undamaged receiver plants exposed to CPVs or HIPVs from *A. argillacea* or *S. littoralis*. Priming effects were tested by testing for effects of treatment (HIPV exposure and /or insect damage) and the interaction with the insect species. Post-hoc tests were performed as tukey-adjusted pairwise comparisons.

| **Phytohormone** | **Test** | **Comparison** | **Test value** | **Df** | **p-value** |
|---|---|---|---|---|---|
| **Testing for induction** | | | | | |
| JA | Anova | Treatment | $\chi^2$ = 54.095 | **2** | < 0.0001 |
| | Post-hoc | CPVs vs. HIPVs (*A. argillacea)* | t = -6.393 | 16 | < 0.0001 |
| | | CPVs vs. HIPVs (*S. littoralis*) | t = -7.718 | 16 | < 0.0001 |
| | | HIPVs (*A. argillacea*) vs. HIPVs (*S. littoralis*) | t = -1.617 | 16 | 0.2674 |
| JA-Ile | Anova | Treatment | $\chi^2$ = 29.789 | **2** | < 0.0001 |
| | Post-hoc | CPVs vs. HIPVs (*A. argillacea)* | t = -4.968 | 16 | 0.0004 |
| | | CPVs vs. HIPVs (*S. littoralis*) | t = -5.087 | 16 | 0.0003 |
| | | HIPVs (*A. argillacea*) vs. HIPVs (*S. littoralis*) | t = -0.311 | 16 | 0.95 |
| **Testing for priming effects** | | | | | |
| JA | Anova | Treatment | F = 2.7681 | 2 | 0.079 |
| | | Insect | F = 3.0506 | 1 | 0.091 |
| | | Treatment:Insect | F = 1.0964 | 2 | 0.34 |
| | Post-hoc | HIPVs-undamaged vs. CPVs-undamaged (*A. argillacea*) | t = 0.238 | 30 | 0.97 |
| | | HIPVs-undamaged vs. HIPVs-damaged (*A. argillacea*) | t = -0.378 | 30 | 0.92 |
| | | CPVs-damaged vs. HIPVs-damaged (*A. argillacea*) | t = -0.584 | 30 | 0.83 |
| | | HIPVs-undamaged vs. CPVs-undamaged (*S. littoralis*) | t = 0.786 | 30 | 0.71 |
| | | HIPVs-undamaged vs. HIPVs-damaged (*S. littoralis*) | t = -1.871 | 30 | 0.16 |
| | | CPVs-damaged vs. HIPVs-damaged (*S. littoralis*) | t = -2.683 | 30 | 0.031 |
| JA-Ile | Anova | Treatment | $\chi^2$ = 10.1506 | 2 | 0.0062 |
| | | Insect | $\chi^2$ = 2.1754 | 1 | 0.14 |
| | | Treatment:Insect | $\chi^2$ = 3.2295 | 2 | 0.20 |
| | Post-hoc | HIPVs-undamaged vs. CPVs-undamaged (*A. argillacea*) | t = -2.090 | 30 | 0.11 |
| | | HIPVs-undamaged vs. HIPVs-damaged (*A. argillacea*) | t = -0.684 | 30 | 0.77 |
| | | CPVs-damaged vs. HIPVs-damaged (*A. argillacea*) | t = 1.259 | 30 | 0.43 |
| | | HIPVs-undamaged vs. CPVs-undamaged (*S. littoralis*) | t = -2.242 | 30 | 0.081 |
| | | HIPVs-undamaged vs. HIPVs-damaged (*S. littoralis*) | t = -2.947 | 30 | 0.016 |
| | | CPVs-damaged vs. HIPVs-damaged (*S. littoralis*) | t = -0.918 | 30 | 0.63 |

**Table S6.** Complete statistics for the phytohormones abscisic acid (ABA), indole-3-acetic acid (IAA), 12-oxophytodienoic acid (OPDA) and salicylic acid (SA). Direct induction was tested by comparing only the undamaged receiver plants exposed to CPVs or HIPVs from *A. argillacea* or *S. littoralis*. Priming effects were tested by testing for effects of treatment (HIPV exposure and /or insect damage) and the interaction with the insect species. Post-hoc tests were performed as tukey-adjusted pairwise comparisons.

| Phytohormone | Test | Comparison | Test value | Df | p-value |
|---|---|---|---|---|---|
| **Testing for induction** | | | | | |
| ABA | Anova | Treatment | $\chi^2$ = 0.48181 | 2 | 0.79 |
| IAA | Anova | Treatment | $\chi^2$ = 0.75647 | 2 | 0.96 |
| OPDA | Anova | Treatment | $\chi^2$ = 4.2161 | 2 | 0.12 |
| SA | Anova | Treatment | $\chi^2$ = 8.5093 | 2 | 0.014 |
| | Post-hoc | CPVs vs. HIPVs (*A. argillacea)* | t = -2.247 | 16 | 0.093 |
| | | CPVs vs. HIPVs (*S. littoralis*) | t = 0.367 | 16 | 0.93 |
| | | HIPVs (*A. argillacea*) vs. HIPVs (*S. littoralis*) | t = 2.627 | 16 | 0.046 |
| **Testing for priming effects** | | | | | |
| ABA | Anova | Treatment | $\chi^2$ = 0.63942 | 2 | 0.73 |
| | | Insect | $\chi^2$ = 0.21512 | 1 | 0.64 |
| | | Treatment:Insect | $\chi^2$ = 2.13535 | 2 | 0.34 |
| IAA | Anova | Treatment | $\chi^2$ = 0.40650 | 2 | 0.82 |
| | | Insect | $\chi^2$ = 0.52353 | 1 | 0.47 |
| | | Treatment: Insect | $\chi^2$ = 2.26380 | 2 | 0.32 |
| OPDA | Anova | Treatment | $\chi^2$ = 1.1512 | 2 | 0.56 |
| | | Insect | $\chi^2$ = 0.0027 | 1 | 0.96 |
| | | Treatment:Insect | $\chi^2$ = 5.8392 | 2 | 0.054 |
| SA | Anova | Treatment | $\chi^2$ = 0.9965 | 2 | 0.61 |
| | | Insect | $\chi^2$ = 4.1312 | 1 | 0.042 |
| | | Treatment:Insect | $\chi^2$ = 4.8724 | 2 | 0.087 |
| | Post-hoc | HIPVs-undamaged vs. CPVs-undamaged (*A. argillacea*) | t = 0.300 | 30 | 0.95 |
| | | HIPVs-undamaged vs. HIPVs-damaged (*A. argillacea*) | t = 2.265 | 30 | 0.076 |
| | | CPVs-damaged vs. HIPVs-damaged (*A. argillacea*) | t = 1.915 | 30 | 0.15 |
| | | HIPVs-undamaged vs. CPVs-undamaged (*S. littoralis*) | t = -0.098 | 30 | 0.99 |
| | | HIPVs-undamaged vs. HIPVs-damaged (*S. littoralis*) | t = -0.697 | 30 | 0.77 |
| | | CPVs-damaged vs. HIPVs-damaged (*S. littoralis*) | t = -0.628 | 30 | 0.81 |

**Table S7.** Complete statistics for the gene expression of the genes *2-ODD and Cdn1c3* as shown in Figure 6A. Direct induction was tested by comparing only the undamaged receiver plants exposed to CPVs or HIPVs from *A. argillacea* or *S. littoralis*. Priming effects were tested by testing for effects of treatment (HIPV exposure and /or insect damage) and the interaction with the insect species. Post-hoc tests were performed as tukey-adjusted pairwise comparisons.

| Gene | Test | Comparison | Test value | Df | p-value |
|---|---|---|---|---|---|
| **Testing for induction** | | | | | |
| *2-ODD* | Anova | Treatment | $\chi^2 = 0.5317$ | **2** | 0.77 |
| *Cdn1c3* | Anova | Treatment | $\chi^2 =$ | **2** | 0.29 |
| **Testing for priming effects** | | | | | |
| *2-ODD* | Anova | Treatment | $\chi^2 = 34.628$ | 2 | < 0.0001 |
| | | Insect | $\chi^2 = 1.454$ | 1 | 0.23 |
| | | Treatment:Insect | $\chi^2 = 3.2295$ | 2 | 0.89 |
| | Post-hoc | HIPVs-undamaged vs. CPVs-undamaged (*A. argillacea*) | t = -4.364 | 30 | 0.0004 |
| | | HIPVs-undamaged vs. HIPVs-damaged (*A. argillacea*) | t = -3.854 | 30 | 0.0016 |
| | | CPVs-damaged vs. HIPVs-damaged (*A. argillacea*) | t = 0.283 | 30 | 0.96 |
| | | HIPVs-undamaged vs. CPVs-undamaged (*S. littoralis*) | t = -4.378 | 30 | 0.0004 |
| | | HIPVs-undamaged vs. HIPVs-damaged (*S. littoralis*) | t = -4.574 | 30 | 0.0002 |
| | | CPVs-damaged vs. HIPVs-damaged (*S. littoralis*) | t = -0.196 | 30 | 0.98 |
| *Cdn1c3* | Anova | Treatment | $\chi^2 = 13.219$ | 2 | 0.0013 |
| | | Insect | $\chi^2 = 0.9378$ | 1 | 0.33 |
| | | Treatment:Insect | $\chi^2 = 0.0685$ | 2 | 0.97 |
| | Post-hoc | HIPVs-undamaged vs. CPVs-undamaged (*A. argillacea*) | t = -2.513 | 30 | 0.045 |
| | | HIPVs-undamaged vs. HIPVs-damaged (*A. argillacea*) | t = -2.791 | 30 | 0.024 |
| | | CPVs-damaged vs. HIPVs-damaged (*A. argillacea*) | t = -0.391 | 30 | 0.92 |
| | | HIPVs-undamaged vs. CPVs-undamaged (*S. littoralis*) | t = -2.149 | 30 | 0.097 |
| | | HIPVs-undamaged vs. HIPVs-damaged (*S. littoralis*) | t = -2.482 | 30 | 0.048 |
| | | CPVs-damaged vs. HIPVs-damaged (*S. littoralis*) | t = -0.333 | 30 | 0.94 |

**Table S8.** Complete statistics for the terpene aldehydes gossypol, hemigossypolone and heliocides as shown in Figure 6B. Direct induction was tested by comparing only the undamaged receiver plants exposed to CPVs or HIPVs from *A. argillacea* or *S. littoralis*. Priming effects were tested by testing for effects of treatment (HIPV exposure and /or insect damage) and the interaction with the insect species. In all models, leaf mass was added as a covariate. Post-hoc tests were performed as tukey-adjusted pairwise comparisons.

| **Terpene aldehyde** | **Test** | **Comparison** | **Test value** | **Df** | **p-value** |
|---|---|---|---|---|---|
| **Testing for induction** | | | | | |
| Gosspyol | Anova | Treatment | $\chi^2 = 0.5897$ | 2 | 0.74 |
| | | 4th leaf mass | $\chi^2 = 14.5580$ | 1 | 0.00014 |
| Hemigossypolone | Anova | Treatment | $\chi^2 = 0.78253$ | 2 | 0.68 |
| | | 4th leaf mass | $\chi^2 = 1.7538$ | 1 | 0.19 |
| Heliocides | Anova | Treatment | $\chi^2 = 5.2995$ | 2 | 0.071 |
| | | 4th leaf mass | $\chi^2 = 3.11590$ | 1 | 0.078 |
| **Testing for priming effects** | | | | | |
| Gossypol | Anova | Treatment | $\chi^2 = 26.915$ | 2 | < 0.0001 |
| | | Insect | $\chi^2 = 0.951$ | 1 | 0.33 |
| | | Treatment:Insect | $\chi^2 = 38.044$ | 2 | < 0.0001 |
| | | 4th leaf mass | $\chi^2 = 5.592$ | 1 | 0.061 |
| | Post-hoc | HIPVs-undamaged vs. CPVs-undamaged (*A. argillacea*) | t = 1.094 | 30 | 0.52 |
| | | HIPVs-undamaged vs. HIPVs-damaged (*A. argillacea*) | t = -2.857 | 30 | 0.020 |
| | | CPVs-damaged vs. HIPVs-damaged (*A. argillacea*) | t = -4.331 | 30 | 0.0004 |
| | | HIPVs-undamaged vs. CPVs-undamaged (*S. littoralis*) | t = -1.931 | 30 | 0.15 |
| | | HIPVs-undamaged vs. HIPVs-damaged (*S. littoralis*) | t = -3.551 | 30 | 0.0033 |
| | | CPVs-damaged vs. HIPVs-damaged (*S. littoralis*) | t = -1.617 | 30 | 0.25 |
| Hemigossypolone | Anova | Treatment | $\chi^2 = 28.4210$ | 2 | < 0.0001 |
| | | Insect | $\chi^2 = 0.2616$ | 1 | 0.61 |
| | | Treatment:Insect | $\chi^2 = 3.6124$ | 2 | 0.16 |
| | | 4th leaf mass | $\chi^2 = 19.7093$ | 1 | < 0.0001 |
| | Post-hoc | HIPVs-undamaged vs. CPVs-undamaged (*A. argillacea*) | t = 0.478 | 30 | 0.88 |
| | | HIPVs-undamaged vs. HIPVs-damaged (*A. argillacea*) | t = -3.111 | 30 | 0.010 |
| | | CPVs-damaged vs. HIPVs-damaged (*A. argillacea*) | t = -3.933 | 30 | 0.0012 |
| | | HIPVs-undamaged vs. CPVs-undamaged (*S. littoralis*) | t = -1.995 | 30 | 0.13 |
| | | HIPVs-undamaged vs. HIPVs-damaged (*S. littoralis*) | t = -3.761 | 30 | 0.0019 |
| | | CPVs-damaged vs. HIPVs-damaged (*S. littoralis*) | t = -1.763 | 30 | 0.20 |
| Heliocides | Anova | Treatment | $\chi^2 = 59.249$ | 2 | < 0.0001 |
| | | Insect | $\chi^2 = 0.068$ | 1 | 0.79 |
| | | Treatment:Insect | $\chi^2 = 4.531$ | 2 | 0.10 |
| | | 4th leaf mass | $\chi^2 = 6.487$ | 1 | 0.011 |
| | Post-hoc | HIPVs-undamaged vs. CPVs-undamaged (*A. argillacea*) | t = 0.545 | 30 | 0.85 |
| | | HIPVs-undamaged vs. HIPVs-damaged (*A. argillacea*) | t = -3.658 | 30 | 0.0025 |
| | | CPVs-damaged vs. HIPVs-damaged (*A. argillacea*) | t = -4.605 | 30 | 0.0002 |
| | | HIPVs-undamaged vs. CPVs-undamaged (*S. littoralis*) | t = -2.453 | 30 | 0.050 |
| | | HIPVs-undamaged vs. HIPVs-damaged (*S. littoralis*) | t = -6.155 | 30 | < 0.0001 |
| | | CPVs-damaged vs. HIPVs-damaged (*S. littoralis*) | t = -3.708 | 30 | 0.0022 |

**Table S9.** Primers used for real-time qPCR.

| GeneID | GenBank Accession Numer or Cottongen ID | Forward primer sequence (5'-…-3') | Reverse primer sequence (5'-…-3') |
|---|---|---|---|
| *GhCDN1-C3* | AF174294 | AACTCAAAAACGCCACCAAC | TAGTCGGAATCGAAGGGATG |
| *Gh2ODD* | Gh_A01G1964 | ATTGGTGCTTGTTTACGGGT | GGAGCAGAGTCAGGCCAC |
| Histone | AF024716 | GAAGCCTCATCGATACCGT | CTACCACTACCATCATGGC |
